# Computation is concentrated in rich clubs of local cortical networks

**DOI:** 10.1101/290981

**Authors:** Samantha P. Faber, Nicholas M. Timme, John M. Beggs, Ehren L. Newman

## Abstract

To understand how neural circuits process information, it is essential to identify the relationship between computation and circuit organization. Rich-clubs, highly interconnected sets of neurons, are known to propagate a disproportionate amount of information within cortical circuits. Here, we test the hypothesis that rich-clubs also perform a disproportionate amount of computation. To do so, we recorded the spiking activity of on average ∼300 well-isolated individual neurons from organotypic cortical cultures. We then constructed weighted, directed networks reflecting the effective connectivity between the neurons. For each neuron, we quantified the amount of computation it performed based on its inputs. We found that rich-club neurons compute ∼160% more information than neurons outside of the rich club. The amount of computation performed in the rich-club was proportional to the amount of information propagation by the same neurons. This suggests that, in these circuits, information propagation drives computation. In total, our findings indicate that rich club organization in effective cortical circuits supports not only information propagation but also neural computation.

**AUTHOR SUMMARY:** Here we answer the question of whether rich club organization in functional networks of cortical circuits supports neural computation. To do so, we combined network analysis with information theoretic tools to analyze the spiking activity of hundreds of neurons recorded from organotypic cultures of mouse somatosensory cortex. We found that neurons in rich clubs computed significantly more than neurons outside of rich clubs, suggesting that rich-clubs do support computation in cortical circuits. Indeed, the amount of computation that we found in the rich clubs was proportional to the amount of information they propagate suggesting that, in these circuits, information propagation drives computation.

## INTRODUCTION

The idea that neurons propagate information and that downstream neurons integrate this information via neural computation is foundational to our understanding of how the brain processes, and responds to, the world. Yet, the determinants of such computations remain largely unknown. Advances in data acquisition methods, offering increasingly comprehensive recordings of the activation dynamics that play out atop of neural circuits, together with advances in data analytics now make it possible to empirically study the determinants of neural computation. Using these tools, we addressed the fundamental question of where, relative to information flow in local cortical networks, the majority of neural computation takes place.

Although there is no agreed upon definition of “computation,” in its simplest form, computation refers to the process of integrating multiple sources of information to produce a new output. This is in contrast to information propagation which simply passes (unmodified) information from a source to a receiver. Neural computation is the systematic transformation of information received by a neuron (determined by analyzing its output with respect to its inputs) based on the input of multiple upstream neurons (Timme et al., 2016). This type of computation can be detected empirically when the activity of upstream neurons accounts for the activity of a downstream neuron better when considered jointly than when treated as independent sources of variance. Because computation is the information gained beyond what was already accounted for by the upstream neurons when they are treated independently, it is not a given that strong sources of information propagation necessarily lead to a large amount of computation. Analytical tools adapted from Shannon’s information theory make it possible to track such computation as well as information propagation in networks of spiking neurons (Strong et al., 1998; Borst & Theunissen, 1999; Schreiber, 2000; Williams & Beer, 2010).

The determinants of strong computation in neural circuits remain poorly understood. Previously, Timme et al. (2016) used these tools to show that computation does not vary systematically with the number of inputs received by a neuron, as might be intuited. Rather, computation correlates with the number of outputs of the upstream neurons. This counter-intuitive finding suggested that the amount of computation a neuron performs may be better predicted by its position in the broader topographic structure of the circuit than by the local connectivity. The relationship between computation and the strength of inputs, however, remains unknown. To understand the determinants of maximal computation in neural circuits, it is important to determine how computation varies as a function of the topology of the functional networks along which information propagates and within which computation is performed. This raises the question of what topological conditions support computation.

Local cortical networks, like many complex networks, contain ‘rich clubs.’ That is, the most strongly connected neurons interconnect with a higher probability than would be expected by chance. The existence of a rich club in a functional network predicts that a select set of highly-integrated nodes handles a disproportionately large amount of traffic. Indeed, in the local networks of cortical circuits, 20% of the neurons account for 70% of the information propagation (Nigam et al., 2016). As such, rich clubs represent a conspicuous topographic landmark in the flow of information across neural circuits.

Thus, here we addressed the critical question: What is the role of the rich club with respect to neural computation? We tested among three possible hypotheses. First, computation is constant throughout the topology of a network–predicting that rich clubs do not perform more or less computation than would be expected by chance. Second, computation grows with increasing information availability–predicting that rich clubs are rich in computation given their high information density. Third, computation decreases with increasing information availability– predicting that rich clubs are computationally poor.

To test these hypotheses, we recorded the spiking activity of hundreds of neurons from organotypic cultures of mouse somatosensory cortex and assessed the distribution of computation inside versus outside of rich clubs of information propagation (Figure 1). The results demonstrate that rich club neurons perform 160% more computation than neurons outside of the rich club, accounting for the majority of network computation (∼88%). Throughout the networks, computation occurs proportionally to information propagation (i.e. the two are correlated) though at a slightly reduced rate (∼3% decrease) inside of rich clubs. Importantly, however, rich clubs contained more computation than would be expected given the correlation between propagation and computation. These results show that rich clubs are computationally dense. Thus, rich clubs support elevated amounts of both computation and propagation relative to the rest of the network.

**Figure 1.**
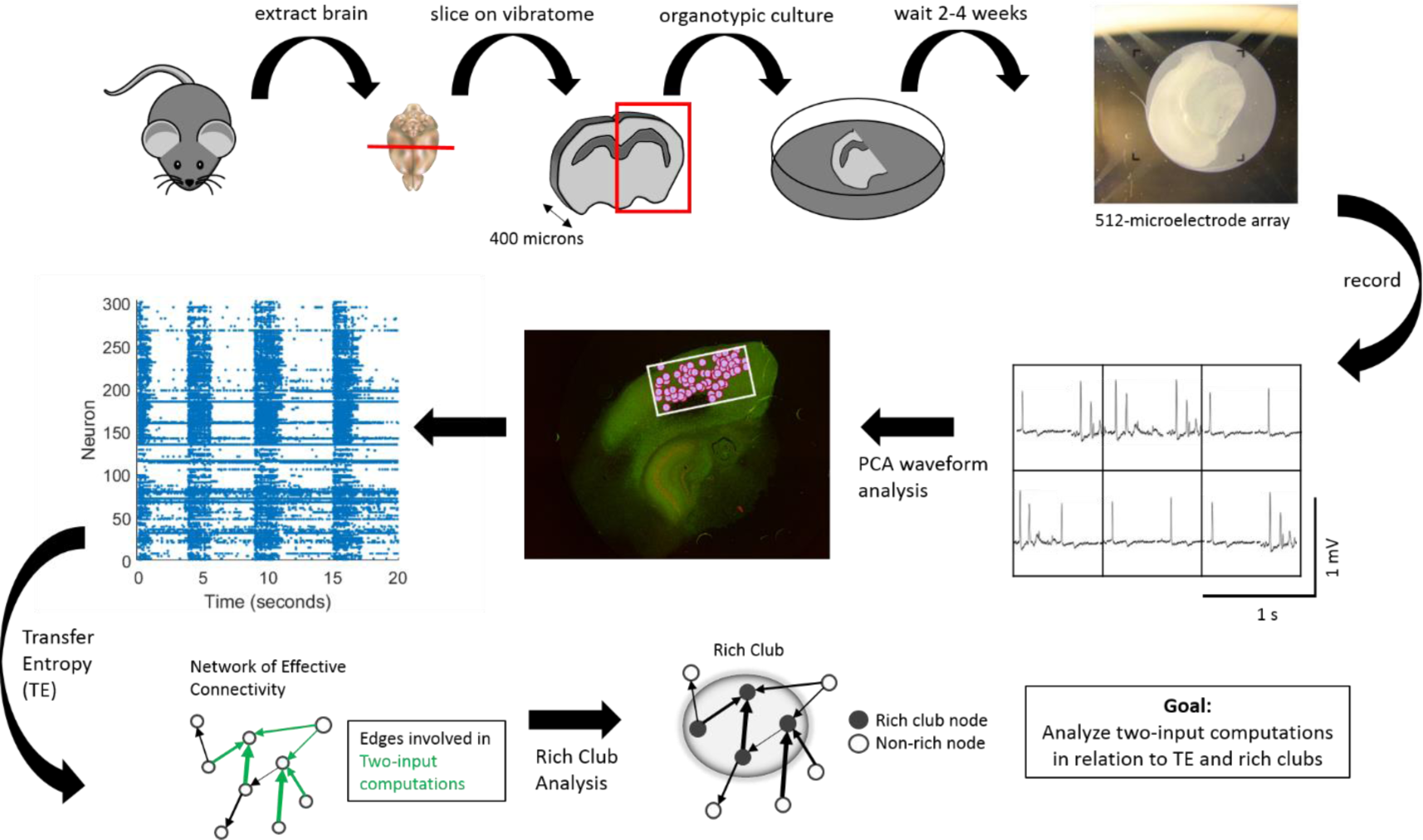
Experimental and data analysis procedure. (Top row, left to right) Brains were extracted from mouse pups and sliced using a vibratome. Slices containing somatosensory cortex were organotypically cultured for up to 2 to 4 weeks. Cultures were then placed on a recording array and recorded for 1 hour. (Middle row, right to left) Recordings yielded neuron spiking dynamics at each electrode – waveforms at six example electrodes shown – which were sorted using PCA in order to isolate individual cells based on their distinct waveforms. Once cells were isolated and localized (pink circles) within the recording area (white rectangle), their corresponding spike trains could be determined. (Bottom Row, left to right) Spike trains were then used to compute Transfer Entropy, at multiple timescales, between each neuron pair in a recording. This resulted in networks of effective connectivity. Computations occurring at neurons receiving two connections were then calculated using partial information decomposition. A rich club analysis was used to detect collections of hub neurons that connect to each other. Finally, we examined the relationship between TE within a triad and two-input computations as well as between two-input computations and rich clubs.

## RESULTS

To study the relationship between information processing and the functional network structure of cortical microcircuits, we recorded spiking activity of between 98 and 594 neurons (median = 310) simultaneously from 25 organotypic cultures of mouse somatosensory cortex. We then quantified the information transfer from each neuron to every other neuron at timescales relevant for synaptic transmission (up to 14 ms; Mason et al., 1991; Swadlow, 1994) using transfer entropy (TE; Schreiber, 2000). TE was selected for its ability to detect nonlinear interactions and deal with discrete data, such as spike trains. The range of 0.05–14 ms was discretized into three subwindows (0.05 – 3 ms, 1.6 – 6.4 ms and 3.5 – 14 ms) to improve the sensitivity to functional interactions across these delays. This resulted in 75 networks (25 recordings at three timescales). Because TE_A→B_ quantifies how much information is gained about whether neuron B will spike in the next moment obtained by knowing whether A is now spiking, it provides a directional, weighted effective connection. Significant TE values (determined by comparing to the distribution of TE values obtained when we shuffled the respective spike trains) were kept and used to build a graph of the circuit’s functional network. The graph was then used to identify all possible triads–in which two transmitting neurons have significant connections to a common receiver neuron. Thus, the same neuron often participated in many triads, the number of which was correlated with the degree of the neuron. Triads were then used to analyze neural computation. Our analysis of neural computation was predicated on the idea that non-linear integration of multiple inputs implements a form of computation (see also Timme et al., 2016). Thus, we quantified neural computation as the additional information regarding the future state of the receiver that was gained beyond what the transmitting neurons offered individually (after accounting for the redundancy between the transmitting neurons and the past state of the receiver itself). We did this using a partial information decomposition approach (PID) (Williams & Beer, 2010) to calculate synergy. PID is currently the only method available that is capable of quantifying the computation that occurs in the multiway interactions of neuronal triads. We focused on triads as the computational unit, rather than higher order configurations, for several reasons. First, they represent the minimal (fundamental) computational unit, on which higher order units depend. Second, they have previously been demonstrated to account for the vast majority of information gained when considering higher order interactions (Timme et al., 2016). And third, calculating synergy for an interaction with even a single additional input (3 inputs total) is much more computationally intensive and does not offer significantly greater explanatory power.

All TE and synergy values were normalized by the entropy of the receiver neuron in order to cast them in terms of the proportion of the receiver neuron’s capacity that is accounted for by the transmitting neuron, or by computation, respectively. Because of this, all TE and synergy values are in terms of bits per bit. All results are reported as medians with 95% bootstrap confidence intervals (computed using 10,000 iterations) presented in brackets. The three timescales at which TE was computed follow the same pattern of results and are therefore combined for ease of presentation; separate results for each timescale are reported in the supplementary material.

### Computation and information propagation vary widely in cortical microcircuits

When building the networks, we found that 0.52% [0.38% 1.1%] of all possible directed connections between neurons were significant at the α = 0.001 level (e.g., 480 of 81510 possible connections, or 0.59%, were significant in a network of 286 neurons). To consider the amount of information that was used in two-input computations, we defined information propagation as the sum of the two inputs (TE values) converging on a neuron. Across neurons, the distribution of propagation values was approximately lognormal (shown in Figure 2), consistent with previously observed distributions of both structural (Song et al., 2005; Lefort et al., 2009; Ikegaya et al., 2012) and functional connectivity (for a review, see Buzsáki & Mizuseki, 2014). This lognormality indicates that there exists a long tail of large propagation values such that a few neurons propagate particularly large amounts of information. Concretely, we found that the strongest 8.5% [8% 9.7%] of neurons propagated as much information as the rest of the neurons combined. Within a network, the difference between the neurons that received the most versus the least information commonly spanned 3.9 [3.6 4.1] orders of magnitude.

**Figure 2.**
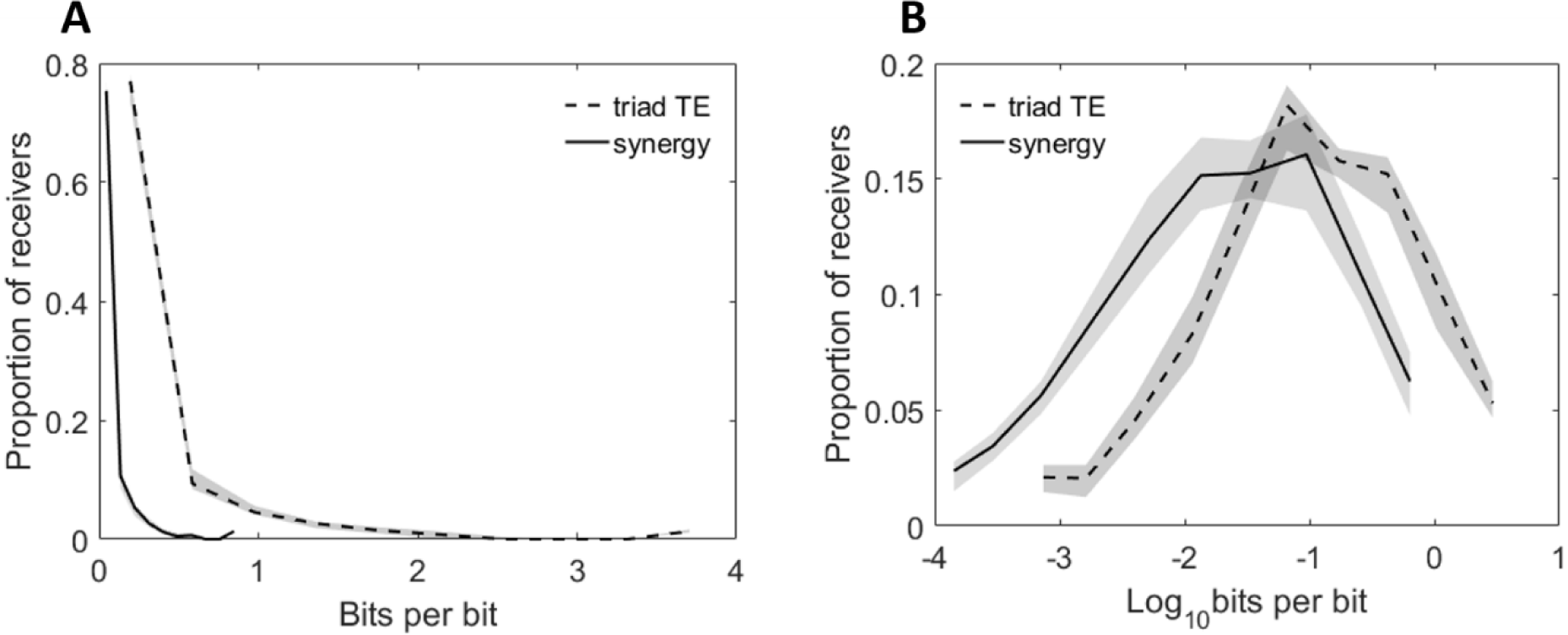
Distributions of neuron computation (synergy) and propagation (triad TE) are highly varied. Histograms of synergy and triad TE values for all receivers in all networks at all timescales. **A.** Distributions of measured values. **B.** Distributions of log-scaled values to emphasize variability. Solid and dashed lines depict the median across networks and shaded regions depict 95% bootstrap confidence intervals around the median.

Computation (measured as synergy), like propagation, varied in a lognormal fashion (Figure 2). The top 8.4% [8% 9.3%] of neurons computed the same amount of information as the rest of the neurons combined. The synergy typically spanned 4.1 [3.8 4.2] orders of magnitude over neurons in individual networks. Notably, computation was reliably smaller than propagation (0.11 vs 0.025; Z_s.r._ = -7.5, n = 75 networks, p = 5.3×10^-14^) indicating that, despite finding substantial computation, most information was accounted for by neuron-to-neuron propagation. The variability in computation motivated us to test the hypothesis that the dense inter-connectivity of rich clubs serves as a hub, not only for propagation, but also for computation.

### Rich clubs perform a majority of the network-wide computation

In every network, we found significant rich clubs at multiple richness parameter (*r*) levels (i.e., thresholds) as shown in Figure 3. The richness parameter was defined as the sum of incoming and outgoing TE edge weights of each neuron. Rich clubs were computed at every *k*^*th*^ value of *r* by dividing the sum of all weights for nodes with *r ≥ r*_*k*_ by the sum of the *n* largest weights in the network, where *n* is the number of edges between neurons with *r ≥ r*_*k*_. The resulting rich club coefficients approach one when the strongest edges connect the neurons in the subsets defined by *r ≥ r*_*k*.._ These rich club coefficients are then normalized by coefficients from null distributions in order to quantify how much rich clubs differ from those expected by chance (see Supplemental Methods for more detail). The more the normalized rich club coefficient diverges from one, the richer the rich club (Figure 3B). When comparing the observed rich club coefficients to those from null distributions, we also calculated p-values to establish significance of the rich clubs. We found significant rich clubs at multiple richness parameter levels, which typically consisted of the top 10% - 50% of the neurons in each network (Figure 3C). The median number of thresholds that resulted in significant rich club coefficients was 4 per network.

**Figure 3.**
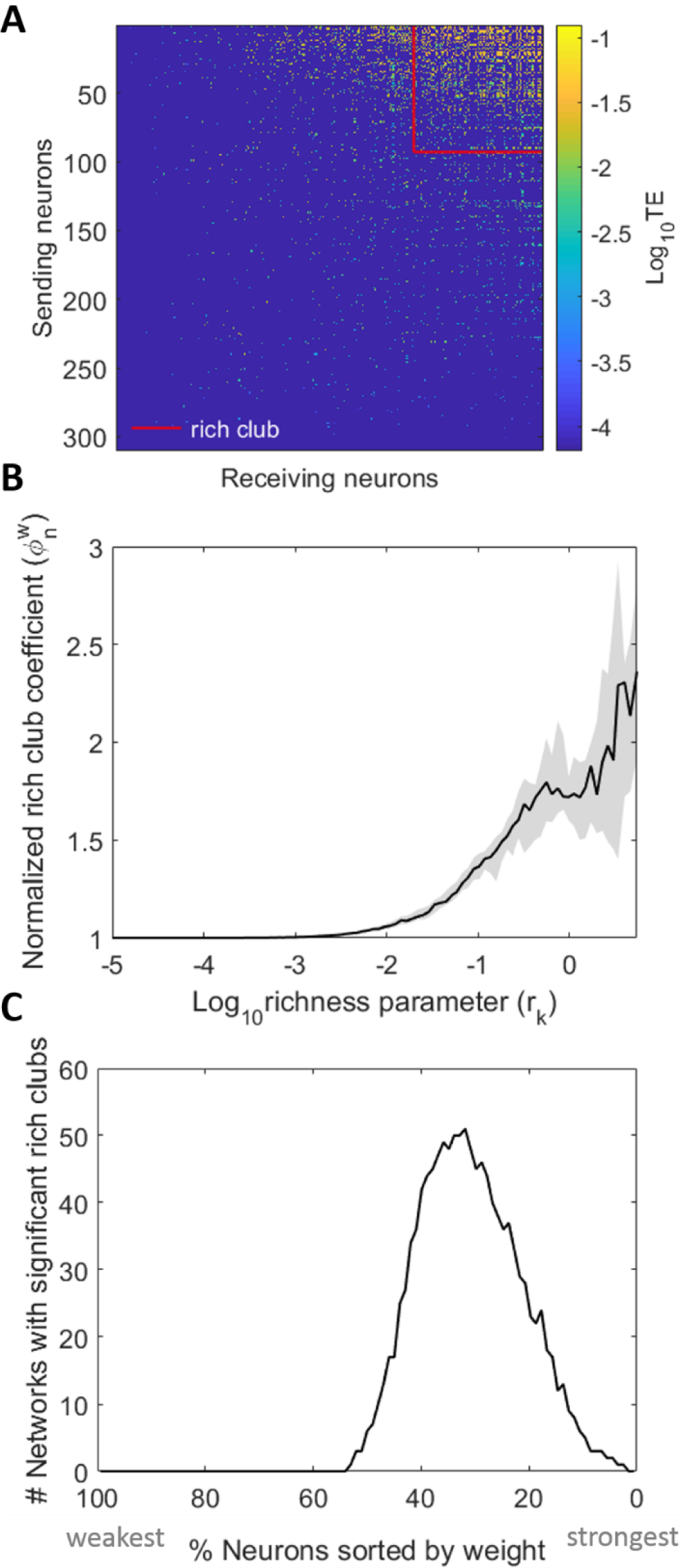
Networks reliably show rich clubs. **A.** Adjacency matrix of a representative 310-neuron network with rich clubs. Rich club of top 30% of neurons depicted. Neurons sorted in order of increasing richness from left to right and bottom to top. TE-values are log-scaled. **B.** Normalized, weighted rich club coefficients for all networks. X-axis is richness parameter level, where the richness parameter is the sum of the weighted connections for each neuron. Solid line represents median across all networks; shaded region is 95% bootstrap confidence interval around the median. In order for a rich club to be recruited into the synergy analysis, coefficients were required to be significant (p < 0.01) when compared to those from randomized networks. **C.** The number of networks, out of the 75 analyzed, with significant rich clubs at each threshold. The majority of networks had significant rich clubs comprised of the top 50% to 10% of the network.

To ensure that the detection of rich clubs was not biased by the spatial sampling of the recording apparatus, we compared the distances between rich club neurons (defined as the top 30% of neurons in a network) to the distances between all neurons in the network. We found that there were no significant differences between the two distributions of distances (KS tests revealed that 75 out of 75 networks had distributions that were not significantly different at the α = 0.01 level).

We investigated the relationship between these rich clubs and computation (measured as synergy) using multiple approaches. In the first approach, we asked if the mean normalized synergy-per-triad (where the mean was taken over all triads in a network, for each network) was significantly greater for triads with receivers inside, versus outside, of a single, representative rich club from each network. The representative rich club for each network was selected at random from among the significant ones. Indeed, we found that mean synergy was 270% greater inside of the rich club (0.027 vs. 0.01, Z_s.r._ = 7.2, n = 75 networks, p = 5.2×10^-13^; Figure 4A). We next asked what percentage of the network-wide computation was performed inside of the rich club and found that, in these representative rich clubs, 87.7% [79.5% 94.3%] of all synergy in a network was performed by rich club neurons (Figure 4B).

**Figure 4.**
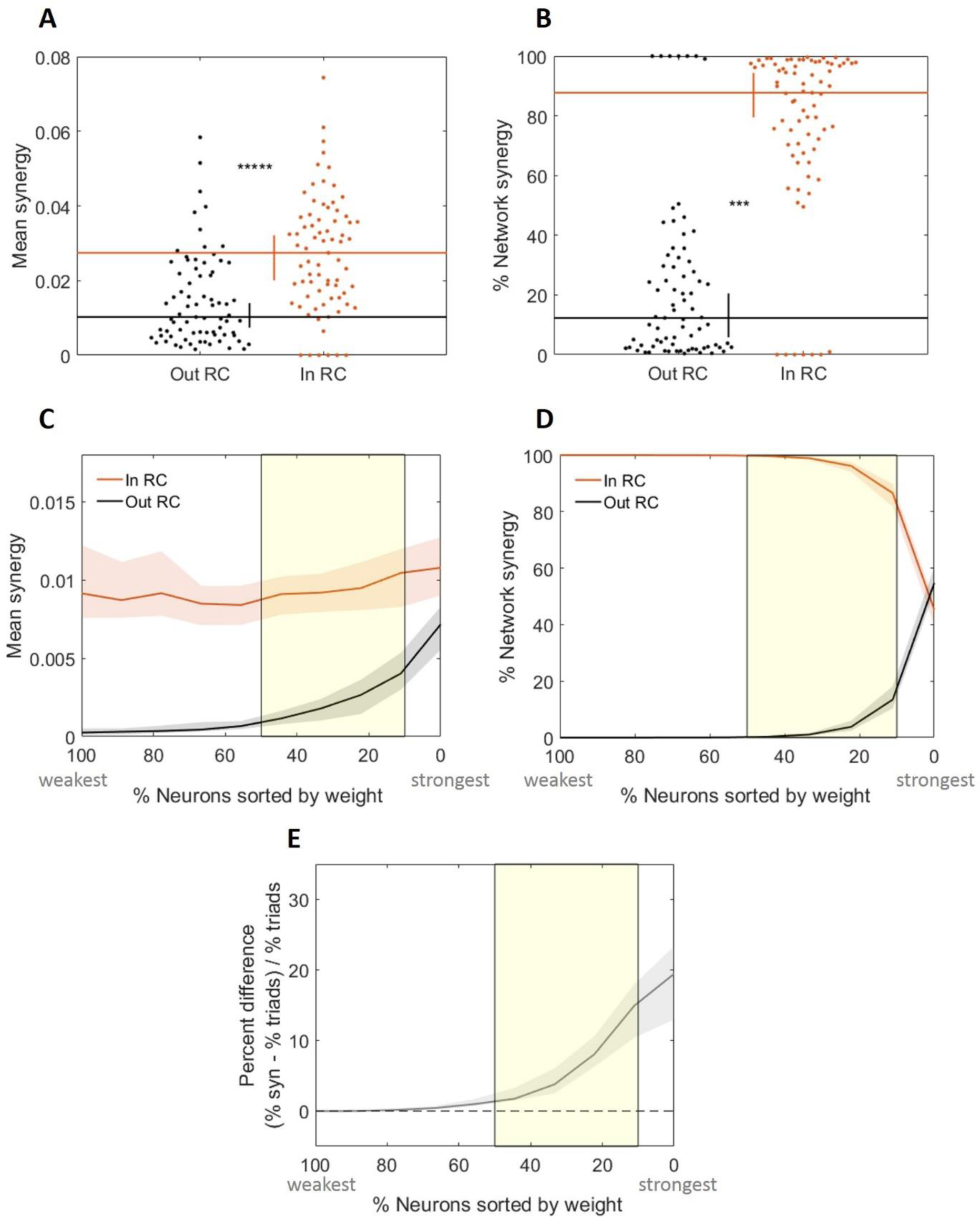
Rich club neurons compute more than non-rich club neurons. **A.** Triads with receivers in the rich club (RC) have greater median mean normalized synergy (compute more) than those with receivers outside the rich club. **B.** Triads with receivers in the RC perform a significantly larger percentage of the total network computation than triads with receivers outside the RC. Distributions shown here are complementary; values sum to 1. **C – E.** Comparison of key metrics at all possible rich club thresholds. The thresholds have been aligned over networks based on the number of neurons in the network that are included at each threshold. The highest (most stringent) thresholds are on the right with the lowest percent of neurons in the rich club. **C.** At all significant rich club levels (indicated by the yellow shaded region), triads with receivers in the rich club have greater median mean synergy than those with receivers outside the rich club. **D.** The percentage of network synergy is plotted as a function of rich club level. At all significant rich club levels, a greater percentage of network synergy occurs in the rich clubs than outside the rich clubs. Distributions shown here are complementary; values sum to 1. **E.** The percent difference in the percentage of network synergy and the percentage of network triads in the rich club is plotted as a function of rich club level. Positive values reflect a larger relative percentage of synergy than percentage of triads. At all significant rich club levels, a greater percentage of synergy is accounted for by a smaller percentage of triads in the rich clubs. Significance indicators: \*\*\*\*\**P* < 1 × 10^-9^; \*\*\**P* < 1 × 10^-6^

A concern that arises when considering whether synergy is stronger inside of rich clubs identified with information theoretic measures, such as TE, is that the rich clubs are defined as having high TE and thus may cause high concentrations of computation for trivial reasons. To test this, we used a spike-time shuffling analysis that preserved TE but disrupted the joint firing statistics between transmitters of a triad. This allowed us to compute the null distribution of synergy that would be expected given the TE that comprised each rich club (see Supplemental Materials for methods). While this analysis demonstrated that high TE in the rich club can be sufficient to generate a null distribution of synergy that is greater inside versus outside of the rich club (significant in two of the three timescales, see supplemental materials for full details), the actual synergy levels observed in the rich club here were reliably even greater than the null distributions. When we Z-scored the observed synergy values by the values of the null distribution of synergy, the median Z-scored synergy values across networks was 13.31 (Z_s.r._ = 5.76, n = 75 networks, p = 8.6×10^-9^). The individual Z-scored synergy values were significant at the α = 0.05 level in 88% (66 of 75) of the networks. The results of these analyses demonstrate that the computation observed in the rich club is not a simple consequence of the magnitude of the TE values that comprise the rich clubs in these networks.

Our second approach to quantify synergy with respect to the rich club examined how synergy varied as a function of the richness levels. The results of these analyses recapitulate the findings reported above. That is, across levels, richer neurons consistently had greater mean synergy-per-triad (Figure 4C). Likewise, at most thresholds, rich neurons accounted for a majority of the network-wide synergy (Figure 4D). However, because the richest neurons are likely to participate in a larger number of triads, rich clubs would be expected to perform a large percentage of the network-wide synergy even if the mean synergy-per-triad was not significantly greater than that found elsewhere in the network. Thus, we also asked how the percentage of network-wide synergy varied as a function of the percentage of triads that are included in the rich club. As shown in Figure 4E, the share of synergy performed by the richest neurons was consistently greater than the percentage of above threshold triads. Notably, the percent difference drops off as the threshold decreases, reflecting that as the rich clubs become less rich, they have less relative amounts of synergy.

Our third approach compared the relationship between the strength of the rich club for any given threshold, as indicated by the normalized rich club coefficient, to the amount of synergy taking place in that rich club. Specifically, we correlated the normalized rich club coefficient with the mean normalized synergy of triads inside of the rich club across thresholds for each network separately and asked if there was a consistent trend across networks (Figure 5). In most networks, synergy and normalized rich club coefficient were positively correlated (64 of 75 networks) such that the median correlation coefficient of 0.75 [0.66 0.84] was significantly greater than zero (Z_s.r._ = 6.6, n = 75 networks, p = 2.9×10^-11^; Figure 5B). These results indicate that rich club strength was strongly predictive of mean synergy.

**Figure 5.**
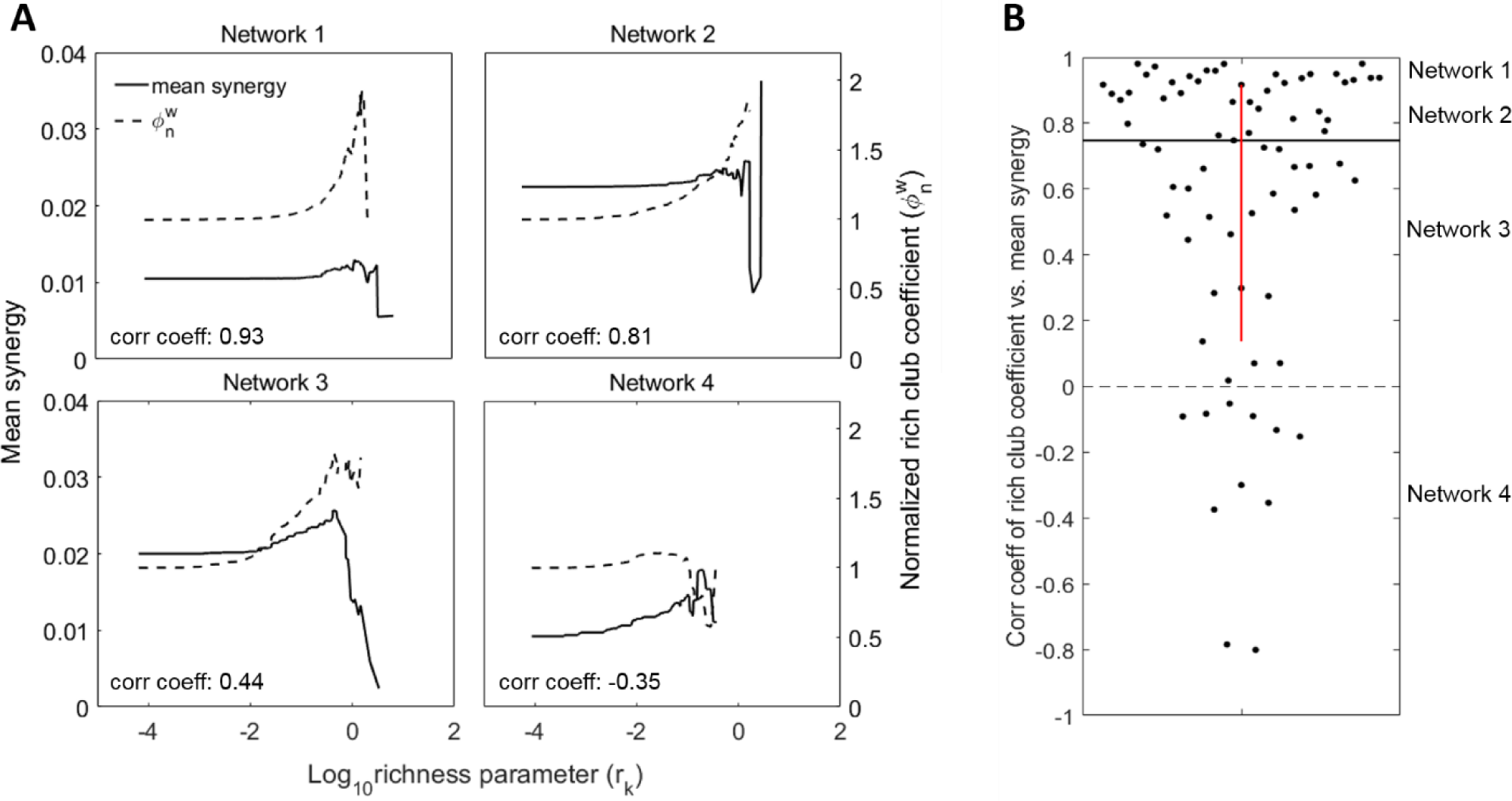
Normalized rich club coefficient correlates with synergy. **A.** Normalized rich club coefficients and mean normalized synergy at increasing richness levels for four representative networks. Negative correlations are observed in networks that have poor rich clubs, or in which the mean synergy decreases as we consider fewer, richer neurons. The second case is observed in networks whose top neurons participate in many triads with synergy values that are highly variable. **B.** Distribution of correlation coefficients for correlations between rich club coefficients and mean triad synergy at all richness levels. Most network rich club coefficients are positively correlated with mean triad synergy. This shows that rich clubs are predictive of increased synergy levels.

Finally, we asked how mean synergy depends upon which triad members were in the rich club. The analyses described above categorize a triad as inside of the rich club if the receiver is a member of the rich club. Here, we calculated the mean normalized synergy separately for each of the six possible alignments of the triad members to the rich club. The results, shown in Figure 6, demonstrate that the single configuration with the largest mean synergy occurred in triads for which all three members were in the rich club. The two configurations with the next greatest mean synergy both had two members in the rich club. Configurations with one member in the rich club had the next greatest mean synergy. Triads that had no neurons in the rich club had the lowest mean synergy. Consistent with the above results, this pattern indicates that the greater the involvement in the rich club, the greater the mean synergy.

**Figure 6.**
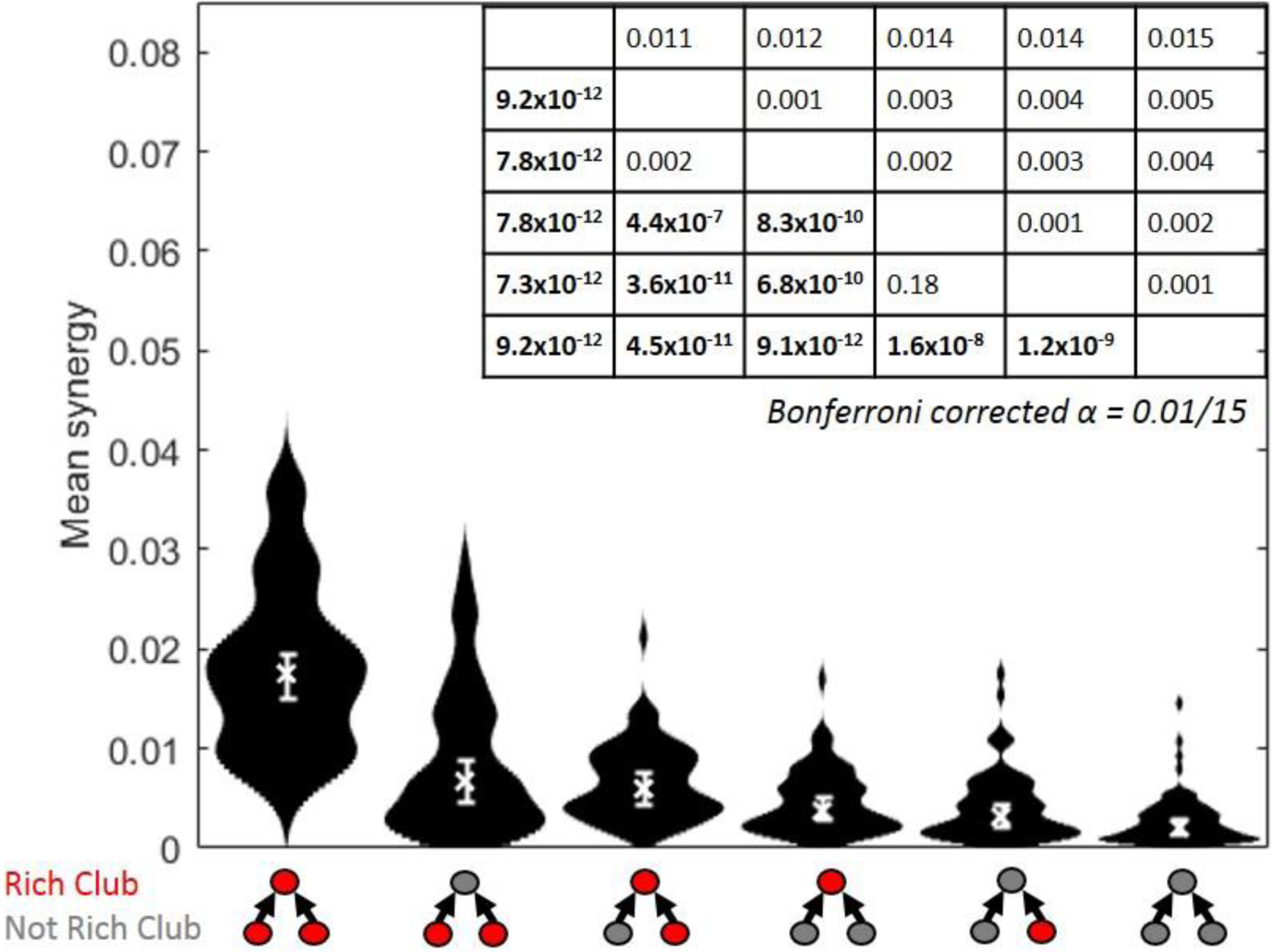
Greater computation (synergy) is performed by triads with greater numbers of neurons in the rich club. Distributions of mean synergy for each of all possible triad interactions with the rich club. Triads that have all members in the rich club have the greatest synergy. Triads with both transmitters in the rich club, and, a single transmitter and the receiver in the rich club have similar amounts of synergy. Triads with only the receiver in the rich club have more synergy than triads with a single transmitter in the rich club. All triads with any member in the rich club have more synergy than triads with no members in the rich club. Medians, denoted by ‘x’, and 95% bootstrap confidence intervals are shown. Table shows Bonferroni-Holm corrected p-values (lower diagonal) and differences of medians (upper diagonal) of pairwise comparisons between the conditions, which are sorted by median mean synergy. Significant p-values are bolded. Distributions shown have n=75 data points.

### Stable computation-to-propagation ratio accounts for the high density of computation in rich clubs

Our results show that rich club neurons both propagate a majority of the information (Nigam et al., 2016) and perform a majority of the computation in the network. A possible explanation for the co-localization of information propagation and computation is that propagation drives computation. To investigate this, we asked how correlated information propagation and computation (measured as synergy) were across triads for each network. In every network, the amount of information being propagated in a triad was strongly correlated with synergy (ρ = 0.76 [0.74 0.77], minimum ρ = 0.57, Z_s.r._ = 7.52, n = 75 networks, p = 5.3×10^-14^; Figures 7A and 7B).

**Figure 7.**
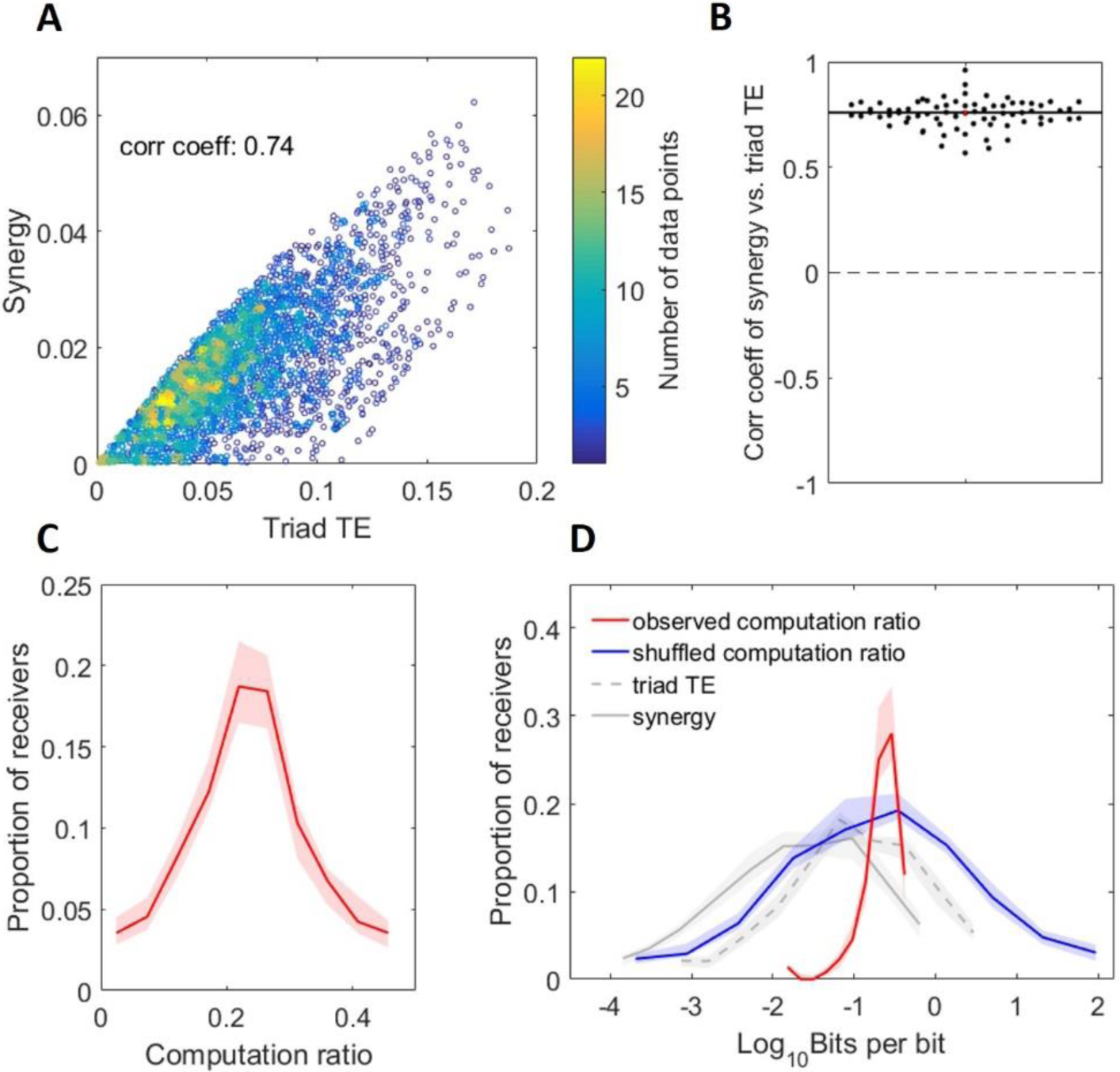
Propagation is highly predictive of computation. **A.** Scatterplot of synergy (computation) versus triad TE (propagation) in a representative network with 3448 triads. Colorbar depicts point density. Also shown is the correlation coefficient. **B.** Distribution of network correlations between synergy and triad TE. This shows that computation was strongly, positively correlated with propagation across all networks. **C.** Histogram of computation ratio values for all receivers in all networks. **D.** Histogram of log-scaled computation ratio values for all receivers in all networks. Gray lines are replotted here from Figure 2 for ease of comparison. The blue line represents the distribution of computation ratios that results from shuffling the alignment of triad synergy to TE. Thus, the span of observed computation ratios is significantly smaller than what we might have observed by chance. For C and D, Solid and dashed lines depict the medians across networks and shaded regions depict 95% bootstrap confidence intervals around the medians.

Seeing that computation and propagation were highly correlated across triads, we asked what range of computation ratios (i.e., computation / propagation) occurred across networks. As shown in Figures 7C and 7D, the computation ratio was highly stereotyped over networks with a median of 0.239 [0.237 0.241]. In contrast to the 3.9 [3.6 4.1] and 4.1 [3.8 4.2] orders of magnitude over which triad TE and synergy varied, respectively, the computation ratio varied by 1.57 [1.46 1.62] orders of magnitude. By way of comparison, randomizing the alignment of synergy to triad TE across triads results in computation ratios that span 5.8 [5.3 6.3] orders of magnitude. The significant reduction in variance over what would be expected by chance (Z_s.r._ =- 7.5, n = 75 networks, p = 5.3×10^-14^) suggests that the computation ratio is a relatively stable property of neural information processing in such networks.

The implication of a stable computation ratio across triads is that information propagation will reliably be accompanied by computation, thereby accounting for the high density of computation in propagation-dense rich clubs. Because the computation ratio allows computation to be predicted from propagation, it is informative to ask if this ratio varies as a function of rich club membership. We found that the computation ratio was not substantially different for triads inside, versus outside, of rich clubs (0.252 vs. 0.259 at the 0.05–3 ms timescale, Z_s.r._= -1.58, n = 25 networks, p = 0.11; 0.238 vs. 0.245 at the 1.6–6.4 ms timescale, Z_s.r._= -2, n = 25 networks, p = 0.045; and 0.215 vs. 0.224 at the 3.5–14 ms timescale, Z_s.r._= 0.97, n = 25 networks, p = 0.33; Figures 8A). The difference was only marginally significant for the 1.6–6.4 ms timescale (Z_s.r._= - 2, n = 25 networks, p = 0.045; Figure 8B, center). We also tested whether the rich club coefficient was correlated with the computation ratio across thresholds and found that it was significantly negatively correlated at the 0.05–3 ms timescale (= -0.36 [-0.69 0.16], Z_s.r._ = -2.38, n = 25 networks, p = 0.017; Figure 8C, left) and at the 1.6–6.4 ms timescale (= -0.62 [-0.70 - 0.18], Z_s.r._ = -3.2, n = 25 networks, p = 0.001; Figure 8C center), but not at the 3.5–14 ms timescale (= -0.26 [-0.69 0.39], Z_s.r._ = -0.98, n = 25 networks, p = 0.33; Figure 8C right). This negative correlation suggest that the computation ratio decreases at high levels of propagation at the shortest timescales. However, computation ratios were only slightly reduced (∼3%) compared to the substantially greater computation values found in the rich club (∼160%), thereby leaving rich clubs overall dense in computation (see Figure 10).

**Figure 8.**
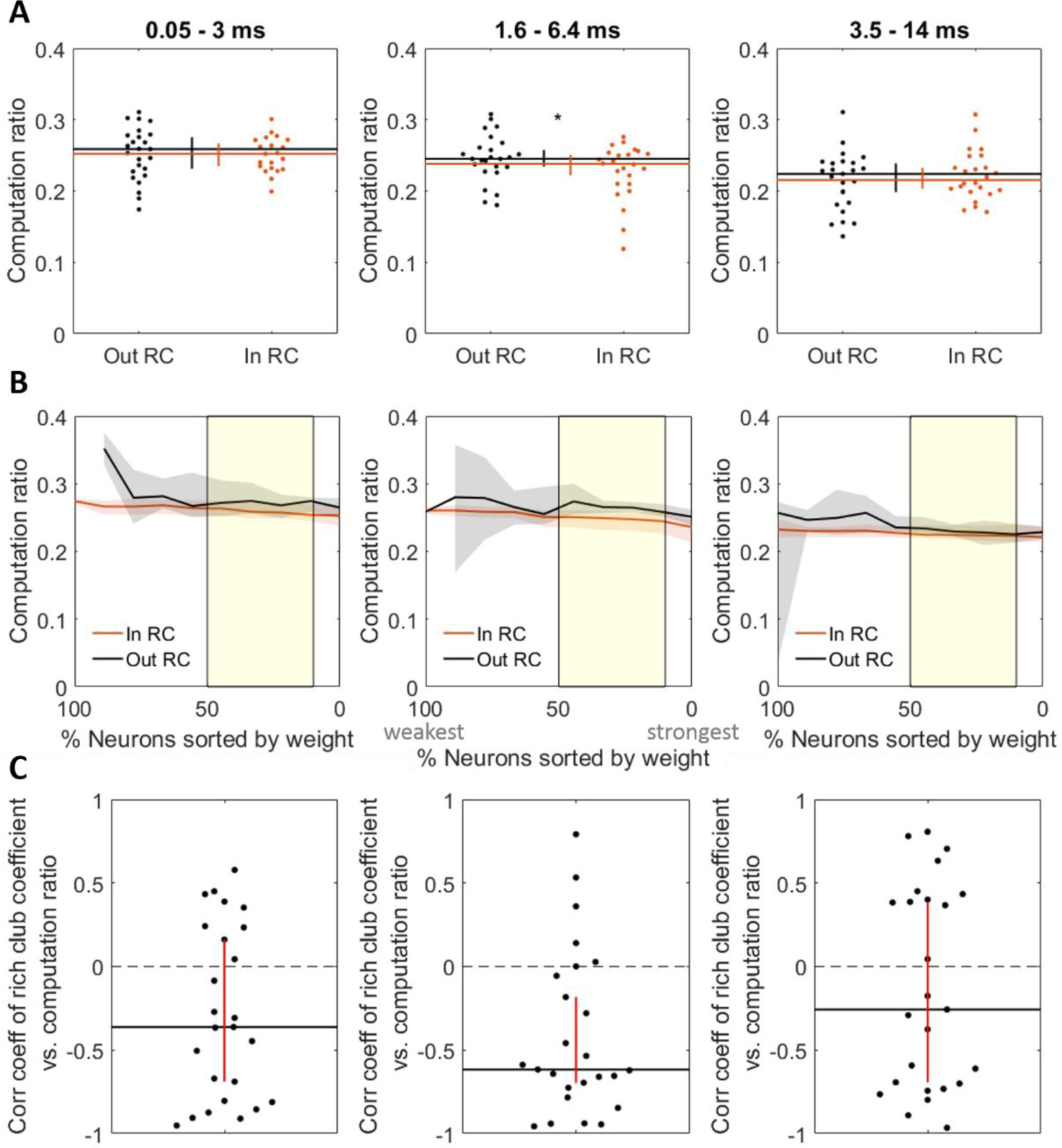
Rich club membership is not strongly predictive of the ratio of computation to propagation (computation ratio). **A**. Median mean computation ratio for triads with receivers inside vs. outside the rich club at the 0.05–3 ms timescale (left), the 1.6–6.4 ms timescale (center), and the 3.5–14 ms timescale (right). **B**. Computation ratio for triads with receivers inside vs. outside the rich club at all significant rich club levels (indicated by the yellow shaded region) at the 0.05–3 ms timescale (left), the 1.6–6.4 ms timescale (center), and the 3.5–14 ms timescale (right). **C**. Coefficient distribution for correlations between mean computation ratio and normalized rich club coefficient at all richness levels, for each network, at the 0.05–3 ms timescale (left), the 1.6–6.4 ms timescale (center), and the 3.5–14 ms timescale (right). Significance indicators: *P < 0.05.

To demonstrate that the result of synergy in the rich club cannot be fully explained by a simple correlation between incoming weight of the receiver and synergy, we asked if there was greater synergy in the rich clubs after the correlation between connection strength and synergy had been regressed out. To do this, we performed a regression between summed incoming connection strength and synergy across triad receivers for a given network and then collected the residual synergy for each triad after accounting for the summed incoming connections. We then asked if the residual synergy values were still significantly greater in the rich club than outside and found that they were (Z_s.r._ = 6.29, n = 75 networks, p = 3.24×10^-10^).

### Operationalization of computation was not critical for present results

To investigate whether our findings are robust to the method of quantifying computation, we implemented two alternate methods of identifying computation to test if the same results were obtained. In the first, we used an alternate implementation of partial information decomposition (PID). Unlike the standard PID approach, which effectively computes the upper bound on synergy (by assuming maximum redundancy between transmitters), this alternate implementation effectively computes the lower bound of synergy (by assuming no redundancy between transmitters). When using this approach, we find the same pattern of results. That is, mean synergy-per-triad is significantly greater inside versus outside of the rich clubs (0.011 vs. 0.006; Z_s.r._ = 5.14, n = 75 networks, p = 2.7×10^-7^; Supplemental Materials Figure 9); and computation is significantly positively correlated with information propagation (ρ = 0.57 [0.42 0.61]; Z_s.r._ = 7.4, n = 75 networks, p = 7.59×10^-14^).

**Figure 9.**
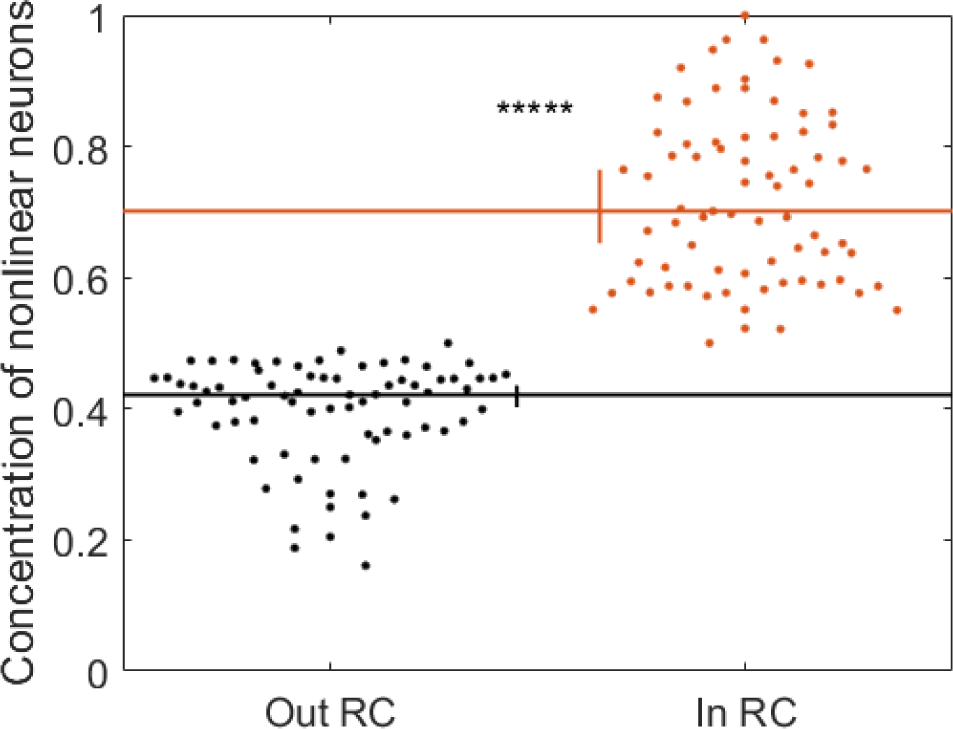
Alternative measure of neural computation reveals results that correspond to those obtained using PID. Neurons with nonlinear transfer functions are represented more inside rich clubs than they are outside rich clubs. Significance indicators: \*\*\*\*\**P* < 1 × 10^-9^

Our second alternate method of identifying computation estimated the input-output transfer functions of individual neurons as described by Chichilnisky (2001). In the prior analyses, using PID, we took synergy as evidence of computation because it quantifies how much more information is carried by multiple neurons when considered together than the sum of information carried by the same neurons when considered individually. Likewise, in this analysis, we took nonlinear input-output functions as evidence of computation because they indicate that a neuron does not simply echo its inputs but, rather, responds based on patterns of upstream inputs. Accordingly, we performed a median split across neurons based on the linearity of their estimated input-output functions. Those with the most nonlinear functions were identified as neurons that likely perform more computation. Comparing this classification to the normalized synergy values obtained via PID, we found that neurons with nonlinear transfer functions were also found to have significantly greater amounts of synergy than those with linear transfer functions (0.109 vs. 0.034; Z_s.r._ = 6.22, n = 75 networks, p = 4.79×10^-10^). As such, use of this classification regime provides an independent, yet related, approach to identifying computation. Consistent with our main results, the concentration of neurons exhibiting nonlinear transfer functions was significantly greater inside versus outside of the rich clubs (70.1% vs. 42.1%; Z_s.r._ = 7.47, n = 75 networks, p = 7.73×10^-14^; Figure 9).

The information theoretic approaches used here to track computation and propagation by / to a receiving neuron are designed to control for the ability of the receiver to account for its own spiking before attributing variance to sending neurons by conditionalizing on the prior state of the receiver. Defining the prior state of the receiver is a parameter dependent process (e.g., the duration and lag of a window in the past to be defined). Here, we used the same window of time to define the past of the receiver as we do the sender. A concern with this approach, however, is that we may be underestimating the information storage of the neuron with use of short windows. To assess the influence this may have on our results, we repeated our analysis with transmitter spike trains for which the timing of each spike was jittered by a random amount drawn from a uniform distribution with a mean of zero and a width of 3 times the duration of the past. This disrupts the short-term interactions but preserves the long-term structure effectively providing a null distribution of values that would be expected given the long-term spiking dynamics alone. By subtracting these values from those obtained from non-jittered spike trains, we were able to assess what effects these values may have had on our findings. We found that, after subtracting these residuals, all results held (Supplemental Figure 11).

## DISCUSSION

Our goal in this work was to test the hypothesis that neural computation in cortical circuits varies in a systematic fashion with respect to the functional organization of those circuits. Specifically, we asked if neurons in rich clubs perform more computation than those outside of the rich clubs. To answer this question, we recorded the spiking activity of hundreds of neurons in organotypic cultures simultaneously and compared the information processing qualities of individual neurons to their relative position in the broader functional network of the circuit. We found that neurons in rich clubs computed ∼160% more than neurons outside of rich clubs. The amount of computation that we found in the rich club was proportional to the amount of information they propagate suggesting that, in these circuits, information propagation drives computation. These results are summarized in Figure 10.

**Figure 10.**
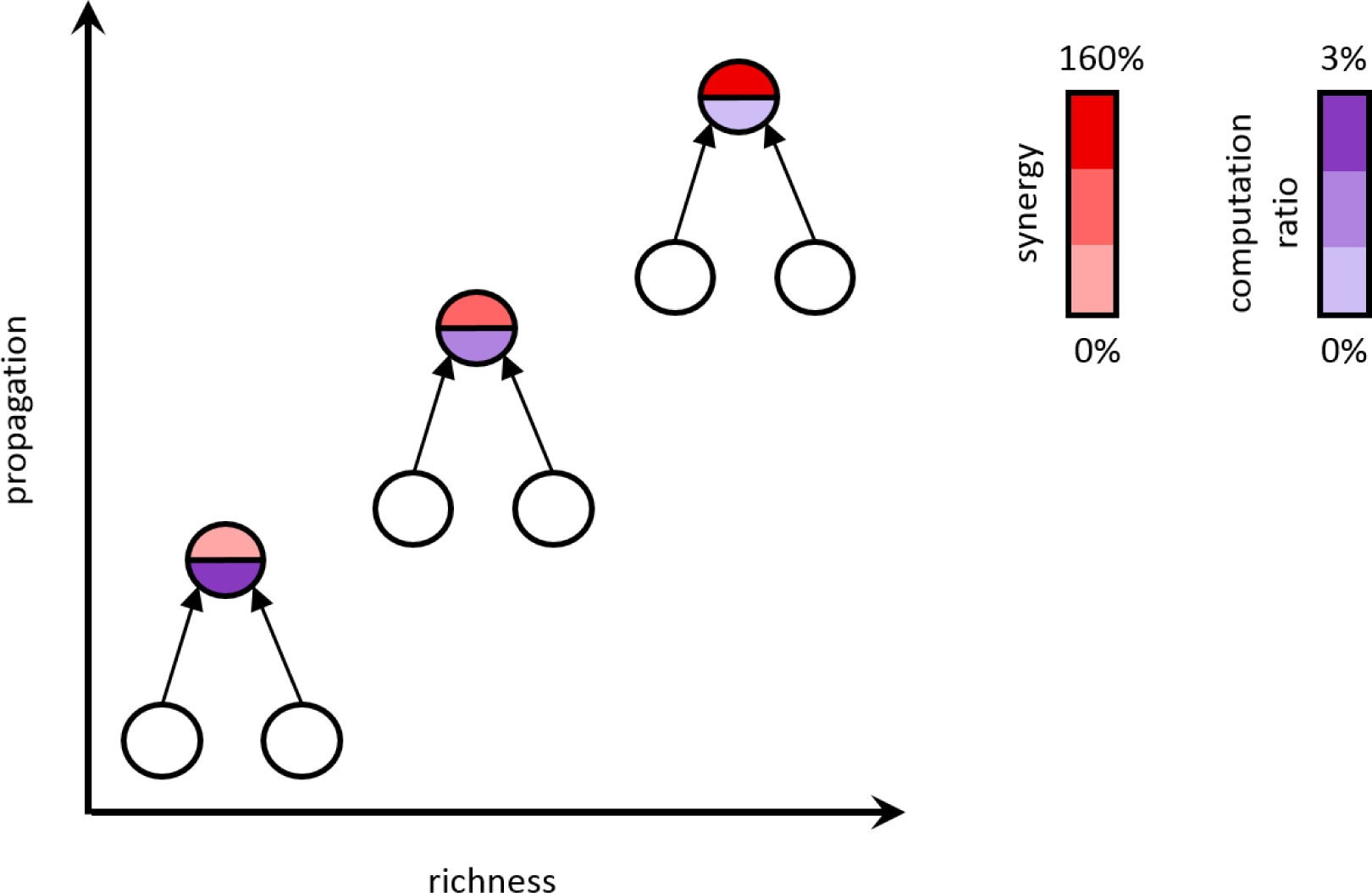
Summary of major findings. Synergy (computation) increases with propagation and node richness by an average of 160% and the computation ratio (amount of computation performed relative to propagation) decreases by 3%.

### Finding computation in organotypic cortical networks

What does it mean for organotypic cultures to process information? In a neural circuit that has no clear sensory inputs, it is easy to imagine that all spiking is either spontaneous or in response to upstream spontaneous spiking. Spontaneous spikes contribute to what, in an information theoretic framework, is considered entropy. When the spiking of upstream neurons allows us to predict future spiking, one colloquially says that those neurons carry information about the future state of the circuit. This is technically accurate because that prediction effectively reduces the uncertainty of the future state of the system. This is what transfer entropy, used in this work, formally measures. The cause of the upstream spiking, whether spontaneous or sensory driven, does not change this logic. As in systems with intact sensory inputs, the neurons in organotypic cultures process information by integrating received synaptic inputs. Though we argue that, in this respect, information processing by organotypic cultures is like the cortical circuits of intact animals, we recognize that important differences accompany the lack of true sensory input. For example, the spatiotemporal structure of the spontaneous spiking that drives activity in cultures is likely fundamentally different from that driven by sensory experience. Understanding how that structure influences information processing will be important to investigate as technologies enabling high temporal resolution recordings of hundreds of neurons *in vivo* mature.

With regard to building an understanding of the drivers of neural computation, the present work substantially builds on previous work that showed that computation was positively correlated with the number of outgoing connections (i.e., out degree) of the upstream neurons (Timme et al., 2016). Though the rich club membership of those neurons was not analyzed, it is reasonable to assume that neurons with high degree would be included in rich clubs. In that respect, our findings are consistent with the previous report. The work of Timme et al. (2016), however, looked only at degree and did not analyze edge weights. Here, we found a strong correlation between synergy and information propagation (i.e., summed edge weights contributing to computation), indicating that computation is strongly dependent upon weight. This substantially alters our understanding of computation as it shows that, beyond the pattern of connections constituting a network, computation is sensitive to the quantity of the information relayed across individual edges.

### Rich clubs as a home for computation

Prior work on rich clubs convincingly argued that rich clubs play a significant role in the routing of network information (van den Heuvel et al., 2011; Harriger et al., 2012; van den Heuvel et al., 2013; Nigam et al., 2016). For example, Nigam et al. (2016) showed, using the same data, that the top 20% richest neurons transmit ∼70% of network information. Here, we showed that information propagation is directly related to computation. Combining this discovery with the previous knowledge that rich clubs perform large amounts of information propagation accounts for the high densities of computation we observed in the cortical circuit rich clubs. Though the precise mechanism of how computation is derived from propagation is unknown, one possibility is that it is the result of what one might consider to be ‘information collisions.’ This idea is based on the finding of Lizier et al. (2010) who demonstrated that the dominant form of information modification (i.e., computation) in cellular automata is the result of collisions between the emergent particles (see also Adamatzky and Durand-Lose, 2012; Bhattacharjee et al., 2016; Sarkar, 2000). In the context of our circuits, the idea is that computation arises when packets of information embedded in the outgoing spike trains of sending neurons collide onto the same receiving neuron in sufficient temporal proximity to alter the way the receiver responds to those inputs. The density of propagating information in rich clubs would proportionately increase the likelihood of such collisions, and thereby increase the amount (and number) of computation(s) performed by the rich club (Flecker et al., 2011).

### Operationalizing information computation and propagation

Our primary analyses used synergy as a proxy for computation among triads consisting of a pair of transmitting neurons and a single receiving neuron, following the methods of Timme et al. (2016). Synergy, as a measure of the information gained when the pair of transmitters is considered jointly over the combined information carried by the neurons individually, provides an intuitively appealing measure of computation (see Timme et al., 2016 for a comprehensive discussion of this relationship). Synergy is a product of partial information decomposition (PID) (Williams & Beer, 2010). However, PID is not the only information theoretic tool available for quantifying neural computation. Our use of PID was motivated by several factors: 1) it can detect linear and nonlinear interactions; 2) it is capable of measuring the amount of information a neuron computes based on simultaneous inputs from other neurons; and 3) it is currently the only method capable of quantifying how much computation occurs in an interaction in which three variables predict a fourth as done here (the future state of the receiver is predicted from the past state of the receiver and two other transmitters).

Concerns have been raised about how PID calculates the redundancy term in that it results in an over-estimation of redundancy and consequently, synergy (Bertschinger et al., 2014; Pica et al., 2017). Here, we addressed this concern by demonstrating that an alternate implementation of PID that minimizes redundancy (and thus synergy) nonetheless yields the same pattern of results. Going further, we also used a non-information theoretic approach to identify neurons that likely perform substantial computation by finding those neurons with nonlinear input-output transfer functions, following the methods of Chichilnisky (2001). This, like our other analyses, showed that the concentration of computation was greater inside of the rich clubs.

Our primary analyses used transfer entropy as a proxy for information propagation. A strength of transfer entropy is that it makes it possible to quantify the mutual information between a sending and receiving neuron after accounting for variance in the receiving neuron spiking that was predictable from its prior state. However, the window used to define the prior state is parameter dependent (e.g., duration, lag from the present). Here, we defined this window so as to match the window that was used to define the sender state (spanning 0.05–3 ms, 1.6–6.4 ms, or 3.5–14 ms for our three timescales, respectively). By doing so, we maintain clear bounds on the timescale at which the functional dynamics were analyzed (selected *a priori* to spantimescales at which synaptic communication occurs). A risk of using this window size, rather than longer, is that it may underestimate the variance in the receiver spiking that can be accounted for by the prior state of the receiver (i.e., without considering the sending neuron) and, thus, result in larger propagation or synergy values. To assess the impact this may have had on our results, following the precedent set by other (Dragoi & Buzsáki, 2006; Nigam et al., 2016) we performed a control analysis in which we jittered the spiking at short timescales and quantified the residual propagation and synergy. By subtracting these residual values from our original (non-jittered) values, we were able to assess what effects these values may have had on our findings. We found that, after subtracting these residuals, all results held.

### Organotypic cultures as a model system

The goal of the present work was to better understand information processing in local cortical networks. To do this, we used a high-density, 512-microelectrode array in combination with organotypic cortical cultures. This approach allowed us to record spiking activity at a temporal resolution (20 kHz; 50 microseconds) that matched typical synaptic delays in cortex (1-3 ms; Mason et al., 1991). The short inter-electrode spacing of 60 microns was within the range of most monosynaptic connections in cortex (Song et al., 2005). This spacing means that the spiking of most cells is picked up by multiple sites and there are few gaps where cells are too far from electrodes to be recorded. The large electrode count allowed us to simultaneously sample hundreds of neurons, revealing complex structures like the rich club. While the cortical layers in organotypic cultures can differ in some respects from those seen *in vivo* (Staal et al., 2011), organotypic cultures nevertheless exhibit very similar synaptic structure and electrophysiological activity to that found *in vivo* (Caeser et al., 1989; Bolz et al., 1990; Götz and Bolz, 1992; Plenz and Aertsen, 1996; Klostermann and Wahle, 1999; Ikegaya et al., 2004; Beggs and Plenz, 2004). The distribution of firing rates in these cultures is lognormal, as seen *in vivo* (Nigam et al., 2016), and the strengths of functional connections are lognormally distributed, similar to the distribution of synaptic strengths observed in patch clamp recordings (Song et al., 2005, reviewed in Buzsáki & Mizuseki, 2014). These features suggested that organotypic cortical cultures serve as a reasonable model system for exploring local cortical networks, while offering an unprecedented combination of large neuron count, high temporal resolution and dense recording sites that cannot currently be matched with *in vivo* preparations.

### Relevance to cognitive health

The increased computation in rich clubs described here may help to explain the functional importance of rich clubs as well as the role of rich clubs in healthy neural functioning. Prior work has shown that cognitively debilitating disorders, including Alzheimer’s disease, epilepsy, and schizophrenia, are associated with diminished rich club organization (van den Heuvel & Sporns, 2011; van den Heuvel et al., 2013; Braun et al., 2015). An implication of our present findings is that such diminished rich club organization would lead to commensurately diminished neural computation. This could account for the impairments observed in such disorders which include loss of memory, consciousness, or mental cohesiveness. Future studies should make use of *in vivo* methods to explore the relationship between computation and behavior.

### Conclusions

The present work demonstrates, for the first time, that synergy is significantly greater inside, versus outside, of rich clubs. Given this, we conclude that rich clubs not only propagate a large percentage of information within cortical circuits, but are also home to a majority of the circuit-wide computation. We also showed that computation was robustly correlated with information propagation, from which, we infer that computation is driven by information availability. Finally, we found that the ratio of computation to propagation was slightly, though significantly, reduced in rich clubs, suggesting that cortical circuits, like human-engineered distributed-computing architectures, may face a communication versus computation trade-off. These results substantially increase what is known regarding computation by cortical circuits.

## MATERIALS AND METHODS

To answer the question of whether rich club neurons perform more computation than do non-rich club neurons in cortical circuits, we combined network analysis with information theoretic tools to analyze the spiking activity of hundreds of neurons recorded from organotypic cultures of mouse somatosensory cortex. Due to space limitations, here we provide an overview of our methods and focus on those steps that are most relevant for interpreting our results. A comprehensive description of all our methods can be found in the Supplemental Materials. All procedures were performed in strict accordance to guidelines from the National Institutes of Health, and approved by the Animal Care and Use Committees of Indiana University and the University of California, Santa Cruz.

### Electrophysiological recordings

All results reported here were derived from the analysis of electrophysiological recordings of 25 organotypic cultures prepared from slices of mouse somatosensory cortex. One hour long recordings were performed at 20 kHz using a 512-channel array of 5 μm diameter electrode and arranged in a triangular lattice with an inter-electrode distance of 60 μm (spanning approximately 0.9 mm by 1.9 mm). Once the data were collected, spikes were sorted using a PCA approach (Ito et al., 2014; Litke et al., 2004; Timme et al., 2014) to form spike trains of between 98 and 594 (median = 310) well isolated individual neurons depending on the recording.

### Network construction

Networks of effective connectivity, representing global activity in recordings, were constructed following Timme et al (2014, 2016). Briefly, weighted effective connections between neurons were established using transfer entropy (TE; Schreiber, 2000). We computed TE at timescales spanning 0.05 – 14 ms to capture neuron interactions at timescales relevant to synaptictransmission. This was discretized into three logarithmically-spaced bins 0.05–3 ms, 1.6–6.4 ms and 3.5–14 ms and separate effective networks were constructed for each timescale resulting in three networks per recording (75 networks total). Only significant TE, determined through comparison to the TE values obtained with jittered spike trains (α = 0.001; 5000 jitters), were used in the construction of the networks. TE values were normalized by the total entropy of the receiving neuron so as to reflect the percentage of the receiver neuron’s capacity that can be accounted for by the transmitting neuron.

### Quantifying computation

Computation was operationalized as synergy, as calculated by the partial information decomposition (PID) approach described by Williams and Beer (2010, 2011). PID compares the measured TE between neurons *TE*(*J* → *I*)and *TE*(*K* → *I*)with the measured multivariate TE between neurons *TE*({*J, K*} → *I*)to estimate terms that reflect the unique information carried by each neuron, the redundancy between neurons, and the synergy (i.e., gain over the sum of the parts) between neurons. Redundancy was computed as per Supplemental equations 8-10.

Synergy was then computed via:

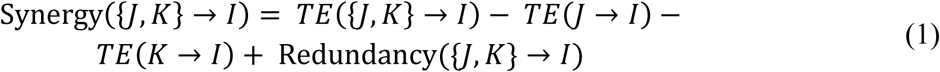

As with TE, synergy was normalized by the total entropy of the receiving neuron. Although there are other methods for calculating synergy (Bertschinger et al., 2014; Pica et al., 2017; Wibral et al., 2017; Lizier et al., 2018), we chose this measure because it is capable of detecting linear and nonlinear interactions and it is currently the only measure which has detailed how one can quantify how much synergy occurs in an interaction in which three variables (here, receiver past and pasts of the two transmitters) predict a fourth (receiver future). Note, we chose not to consider higher order synergy terms, for systems with more than two transmitting neurons, due to the increased computational burden it presented (the number of PID terms increases rapidly as the number of variables increases). However, based on bounds calculated for the highest order synergy term by Timme et al. (2016), it was determined that the information gained by including an additional input beyond two either remained constant or decreased. Thus, it was inferred that lower order (two-input) computations dominated.

### Alternate methods of quantifying computation

To establish that our results are not unique to our approach for quantifying computation, we implemented two alternate methods. The first also uses PID but sets redundancy to be the smallest possible value. Effectively, in this approach synergy is computed as follows:

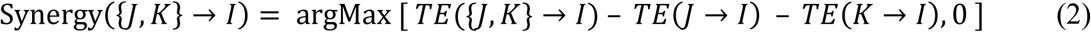

Consequently, synergy is minimized or set to zero when the sum of *TE*(*J*→*I*)and *TE*(*K* →*I*) is greater than *TE*({*J, K*} → *I*). The second alternate method identifies neurons that perform the most computation as those with non-linear input-output transfer functions. The transfer functions are calculated following the methods of Chichilnisky (2001). Briefly, for each neuron, the pattern of inputs across neurons and time that drive the neuron to spike were estimated using a spike triggered average (STA) of the state of all neurons over a 14 ms window prior to each spike. The strength of input to that neuron over time was then estimated by convolving the STA with the time varying state of the network. Finally, the input-output transfer function was established by computing the probability that the neuron fired at each level of input. We then fit both a line and a sigmoid to the resulting transfer function to extract the summed squared error (SSE) that results from each. We categorized neurons with the lowest SSE from the sigmoid fit relative to the SSE from the linear fit as the neurons that perform the most computation.

### Rich club analyses

Weighted rich clubs were identified using a modified version of the rich_club_wd.m function from the Matlab Brain Connectivity toolbox (Rubinov and Sporns, 2010; van den Heuvel and Sporns, 2011), adapted according to Opsahl et al. (2008) to compute weighted rich clubs. To establish the significance of a rich club at a given threshold, we computed the ratio between the observed rich club coefficient and the distribution of those observed when the edges of the network were shuffled. Shuffling was performed according to the methods of Maslov and Sneppen (2002).

To test if synergy was greater for rich club neurons, our first approach randomly selected one of the thresholds–at which the rich club was identified as significant–separately for each network and considered all neurons above that threshold to be in the rich club. Our second approach swept across all possible thresholds, irrespective of the significance of the associated rich club, to assess the influence of varying the ‘richness’ of the neurons treated as if in the rich club. The third approach asked if the results of the second approach were correlated with the strength of the rich club, as quantified via the normalized rich club coefficient, to assess if the strength of the observed effects varied with how much stronger the rich club was than expected by chance.

### Statistics

All results are reported as medians followed by the 95% bootstrap confidence limits (computed using 10,000 iterations), reported inside of square brackets. Likewise, the median is indicated in figures and the vertical bar reflects the 95% bootstrap confidence limits. Comparisons between conditions or against null models were performed using the nonparametric Wilcoxon signed-rank test, unless specified otherwise. The threshold for significance was set at 0.05, unless indicated otherwise in the text.

## ACKNOWLEDGEMENTS

We thank Olaf Sporns and Randy Beer for helpful comments and discussion. This work was sponsored in part by the Whitehall Foundation (PI: EN), MRI instrument grant (NSF number 1429500, PI: JMB), the Robust Intelligence grant (NSF number 1513779, PI: JMB), and the NRT-Interdisciplinary Training in Complex Networks and Systems (NSF number 1735095).

## REFERENCES

Adamatzky A, Durand-Lose J (2012) Collision-Based Computing. In: Handbook of Natural Computing (Rozenberg G, Bäck T, Kok JN, ed). pp. 1949–1978. Berlin, Heidelberg: Springer

Beggs JM, Plenz D (2004) Neuronal avalanches are diverse and precise activity patterns that are stable for many hours in cortical slice cultures. J Neurosci 24:5216 –5229

Bertschinger N, Rauh J, Olbrich E, Jost J, Ay N (2014) Quantifying unique information. Entropy, 16(4), 2161–2183.

Bhattacharjee K, Naskar N, Roy S, Das S (2016) A Survey of Cellular Automata: Types, Dynamics, Non-uniformity and Applications. arXiv:1607.02291

Bolz J, Novak N, Götz M, Bonhoeffer T (1990) Formation of target-specific neuronal projections in organotypic slice cultures from rat visual cortex. Nature 346:359 –362

Borst A, Theunissen F (1999) Information theory and neural coding. Nature Neuroscience 2:947–957

Braun U, Muldoon SF, Bassett DS (2015) On human brain networks in health and disease. eLS.

Buzsáki G, Mizuseki K (2014) The log-dynamic brain: how skewed distributions affect network operations. Nat Rev Neurosci 15:264–278.

Caeser M, Bonhoeffer T, Bolz J (1989) Cellular organization and development of slice cultures from rat visual cortex. Exp Brain Res 77:234 –244.

Chichilnisky EJ (2001) A simple white noise analysis of neuronal light responses. Network: Computation in Neural Systems, 12(2), 199–213.

Dragoi G, Buzsáki, G (2006) Temporal encoding of place sequences by hippocampal cell assemblies. Neuron, 50(1), 145–157.

Flecker B, Alford W, Beggs JM, Williams PL, Beer RD (2011) Partial information decomposition as a spatiotemporal filter. Chaos: An Interdisciplinary Journal of Nonlinear Science 21, 3:037104.

Götz M, Bolz J (1992) Formation and preservation of cortical layers in slice cultures. Journal of Neurobiology. 23: 783–802.

Harriger L, van den Heuvel MP, Sporns O (2012) Rich Club Organization of Macaque Cerebral Cortex and Its Role in Network Communication. PLoS ONE 7(9): e46497.

Ikegaya Y, Aaron G, Cossart R, Aronov D, Lampl I, Ferster D, Yuste R (2004) Synfire chains and cortical songs: temporal modules of cortical activity. Science 304:559 –564

Ikegaya Y, Sasaki T, Ishikawa D, Honma N, Tao K, et al. (2012) Interpyramid spike transmission stabilizes the sparseness of recurrent network activity. Cerebral Cortex 23: 293–304.

Ito S, Yeh FC, Hiolski E, Rydygier P, Gunning DE, Hottowy P, Timme N, Litke AM, Beggs JM (2014) Large-scale, high-resolution multielectrode-array recording depicts functional network differences of cortical and hippocampal cultures. PLoS One 9:e105324.

Klostermann O, Wahle P (1999) Patterns of spontaneous activity and morphology of interneuron types in organotypic cortex and thalamus-cortex cultures. Neuroscience 92: 1243–1259.

Lefort S, Tomm C, Floyd Sarria JC, Petersen CCH (2009) The excitatory neuronal network of the C2 barrel column in mouse primary somatosensory cortex. Neuron 61: 301–316.

Litke A, Bezayiff N, Chichilnisky E, Cunningham W, Dabrowski W, Grillo A, Grivich M, Grybos P, Hottowy P, Kachiguine S (2004) What does the eye tell the brain? Development of a system for the large-scale recording of retinal output activity. IEEE Trans Nucl Sci 51:1434–1440.

Lizier JT, Prokopenko M, Zomaya AY (2010) Information modifications and particle collisions in distributed computation. Chaos: An Interdisciplinary Journal of Nonlinear Science 20(3)

Lizier JT, Bertschinger N, Jost J, Wibral M (2018) Information Decomposition of Target Effects from Multi-Source Interactions: Perspectives on Previous, Current and Future Work. Entropy, 20(4), 307.

Maslov S, Sneppen K (2002) Specificity and Stability in Topology of Protein Networks. Science 03:296, pp. 910–913.

Mason A, Nicoll A, Stratford K (1991) Synaptic transmission between individual pyramidal neurons of the rat visual cortex in vitro. J Neurosci 11:72–84.

Nigam S, Shimono M, Ito S, Yeh FC, Timme N, Myroshnychenko M, Lapish CC, Tosi Z, Hottowy P, Smith WC, Masmanidis SC, Litke AM, Sporns O, Beggs, JM (2016) Rich-Club Organization in Effective Connectivity among Cortical Neurons. J Neurosci 36(4):670–684

Opsahl T, Colizza V, Panzarasa P, Ramasco JJ (2008) Prominence and control: the weighted rich-club effect. Phys Rev Lett 101:168702.

Pica G, Piasini E, Chicharro D, Panzeri S (2017) Invariant components of synergy, redundancy, and unique information among three variables. Entropy, 19(9), 451.

Plenz D, Aertsen A (1996) Neural dynamics in cortex-striatum co-cultures—II. Spatiotemporal characteristics of neuronal activity. Neuroscience 70:893–924.

Rubinov M, Sporns O (2010) Complex network measures of brain connectivity: Uses and interpretations. NeuroImage. 52:1059–69.

Sarkar P (2000) A brief history of cellular automata. ACM Comput Surv 32(1):80–107

Schreiber T (2000) Measuring information transfer. Phys Rev Lett 85:461–464

Song S, Sjöström PJ, Reigl M, Nelson S, Chklovskii DB (2005) Highly non-random features of synaptic connectivity in local cortical circuits. PLoS Biol 3:e68.

Staal JA, Alexander SR, Liu Y, Dickson TD, Vickers JC (2011) Characterization of cortical neuronal and glial alterations during culture of organotypic whole brain slices from neonatal and mature mice. PLOS One 6: e22040

Strong SP, Koberle R, de Ruyter van Steveninck RR, Bialek W (1998) Entropy and information in neural spike trains. Physical Review Letters. 80(1):197–200.

Swadlow HA (1994) Efferent neurons and suspected interneurons in motor cortex of the awake rabbit: axonal properties, sensory receptive fields, and subthreshold synaptic inputs. J Neurophysiol 71:437–453

Tang A, Jackson D, Hobbs J, Chen W, Smith JL, Patel H, Prieto A, Petrusca D, Grivich MI, Sher A, Hottowy P, Dabrowski W, Litke AM, Beggs JM (2008) A maximum entropy model applied to spatial and temporal correlations from cortical networks in vitro. J Neurosci 28:505–518.

Timme NM, Ito S, Myroshnychenko M, Yeh FC, Hiolski E, Litke AM, Beggs JM (2014) Multiplex networks of cortical and hippocampal neurons revealed at different timescales. PLoS One 9: e115764.

Timme NM, Ito S, Myroshnychenko M, Nigam S, Shimono M, Yeh F-C, Hottowy P, Litke AM, Beggs JM (2016) High-Degree Neurons Feed Cortical Computations. PLoS Comput Biol 12(5): e1004858.

van den Heuvel MP, Sporns O (2011) Rich-club organization of the human connectome. J Neurosci 31(44):15775–86.

van den Heuvel MP, Sporns O, Collin G, Scheewe T, Mandl RC, Cahn W,… & Kahn RS (2013) Abnormal rich club organization and functional brain dynamics in schizophrenia. JAMA psychiatry, 70(8), 783–792.

Wibral M, Priesemann V, Kay JW, Lizier JT, Phillips WA (2017) Partial information decomposition as a unified approach to the specification of neural goal functions. Brain and cognition, 112, 25–38.

Williams PL, Beer RD (2010) Nonnegative decomposition of multivariate information. arXiv: 1004.2515.

Williams PL, Beer RD (2011) Generalized measures of information transfer. arXiv: 1102.1507.

