## Supplementary Materials for "Computation is concentrated in rich clubs of local cortical networks" (from another version of this preprint: version 2, 2018-03-29)

### SUPPLEMENTAL METHODS

The methods used in this study include many steps. These will be described below in the following order: (1) in vitro data collection; (2) construction of effective connectivity networks; (3) quantification of neural computation; (4) rich club detection in networks of effective connectivity; (5) quantification of the relationship between synergy and rich clubs; and (6) alternative approaches for quantifying neural computation.

#### *In vitro* data

To study the relationship between neural computation and topological measures of networks of spiking neurons, we analyzed data collected *in vitro*. Data were spontaneously spiking organotypic cultures of mouse somatosensory cortex obtained from postnatal Day 6 to 7 Black 6 mouse pups (RRID:Charles_River:24101632, Harlan) according to Tang et al., 2008 (Ito et al., 2014). All animal tissue samples were prepared according to guidelines from the National Institutes of Health and all animal use procedures were approved by the Indiana University Animal Care and Use Committee as well as the Animal Care and Use Committee at the University of California, Santa Cruz. Spontaneous (as opposed to stimulus-driven) spiking activity in the cultures was recorded at a high temporal resolution of 50 μs, between 2 and 4 weeks after culture preparation, using a 512-microelectrode array (Litke et al., 2004). Array electrodes were flat, 5 μm in diameter and arranged in a triangular lattice with an interelectrode distance of 60 μm. The arrangement allowed for a total recording area of approximately 0.9 mm by 1.9 mm. This preparation and recording method enabled the isolation of large numbers of neurons (an average of 309 cells per recording in 25 hour-long recordings) at high temporal resolution, beyond what can currently be done in any *in vivo* setup. Crucially, the temporal resolution of this method was small enough to resolve synaptic delays of 1-20 ms typically found in cortex (Mason et al., 1991; Swadlow, 1994).

Once the data were collected, spikes were sorted using a PCA approach based on waveforms detected at seven adjacent electrodes (Ito et al., 2014; Litke et al., 2004; Timme et al., 2014). This process yielded a single set of spike times for each isolated neuron. Neurons that spiked fewer than 100 spikes during the hour long recording were removed from the analysis. Spike trains were then used to build networks.

#### Effective connectivity network construction

Because neural computation is fundamentally a dynamic process, we focused on examining networks of effective connectivity. In these networks, connections represent a predictive relationship between the firing of two different neurons. Note, effective connectivity differs from structural connectivity (synapses or gap junctions between neurons) and functional connectivity (e.g., cross-correlations between neuronal time series). Here, effective connections represent directed information transfer between neurons.

Networks of effective connectivity, representing global activity in recordings, were constructed according to Timme et al (2014, 2016) using a measure from information theory known as transfer entropy (TE; Schreiber, 2000). TE was selected for its ability to detect nonlinear interactions and deal with discrete data, such as spike trains. To capture neuron interactions at timescales relevant to synaptic transmission, spiking data was binned at two logarithmically-spaced bin sizes (1.6 and

3.5 ms) and TE was computed at delays (1-4 bins) corresponding to synaptic delays, as in Timme et al. (2014, 2016). Thus, we computed TE at two timescales, 1.6–6.4 ms and 3.5–14 ms. Timescales were purposefully designed to be overlapping so that no interactions were neglected. See Figure 11 for an overview of the binning structure used in TE calculations.

TE quantifies an effective connection from neuron J to neuron I by measuring how much information the past state of the neuron J time series ($J_{t-1}$) produces regarding the current state of the neuron I time series ($I_t$), beyond what is provided by the past state of the neuron I time series ($I_{t-1}$). Here, time series are binary spike trains for neurons I and J, containing 0 for time bins in which the neuron did not spike and 1 for time bins in which it did spike. Generally, the TE from neuron J to neuron I is computed as:

$$TE_{J \to I} = \sum_{i_t,\, i_{t-1},\, j_{t-1}} p(i_t, i_{t-1}, j_{t-1}) \log\left(\frac{p(i_t|\, i_{t-1}, j_{t-1})}{p(i_t|i_{t-1})}\right) \quad (1)$$

The probabilities in Eqn. 1 are computed by counting the number of occurrences of all possible combinations of spiking and not spiking in the $i_t, i_{t-1}$ and $j_{t-1}$ time bins (of the $I_t$, $I_{t-1}$ and $J_{t-1}$ time series) for all bins making up the hour-long recording.

Because we wanted to consider interactions at two timescales associated with synaptic transmission, we included a delay between the past and present states of the neurons so that $i_{t-1}$ became $i_{t-d}$ and $j_{t-1}$ became $j_{t-d}$. Additionally, in order to ensure overlapping timescales, we combined the $i_{t-d}$ and $j_{t-d}$ bins with their previous time bins, such that a spike in either or both time bins corresponded to a state of 1 while no spikes in either time bin corresponded to a state of 0 (See Figure 11 for binning structure). Denoting these new bins as $i'_{t-d}$ and $j'_{t-d}$ gives a slightly different form for TE:

$$TE(d)_{J \to I} = \sum_{i_t,\, i'_{t-d},\, j'_{t-d}} p(i_t,\, i'_{t-d},\, j'_{t-d}) \log\left(\frac{p(i_t|\, i'_{t-d},\, j'_{t-d})}{p(i_t|\, i'_{t-d})}\right) \quad (2)$$

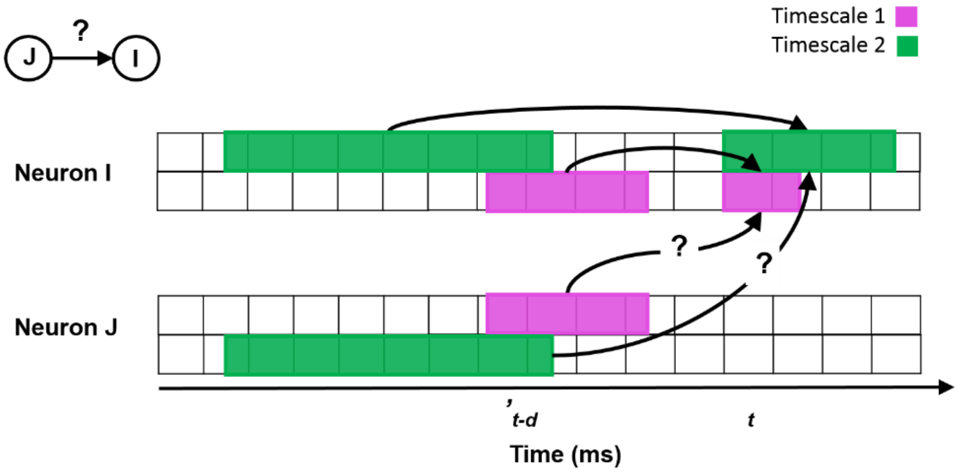

**Figure 11.** *Overview of time series binning structure used in transfer entropy calculations.* Transfer entropy was used to quantify a directed, functional connection from neuron J to neuron I which represents how well the current state (t) of neuron I can be predicted by the past state ('t-d) of neuron J, beyond what is known from the past state of neuron I itself. Two timescales were considered, each with corresponding delays (d). Timescale 1 considered transfer entropy from 1.6—6.4 ms and timescale 2 considered transfer entropy from 3.5—14 ms.

To cast TE in terms of the percentage of the receiver neuron's capacity that can be accounted for by the transmitting neuron, rather than it representing the amount of information being transmitted from transmitter to receiver, we normalized TE by the entropy of the receiver neuron via:

$$TE_{Norm}(d)_{J\to I} = \frac{TE(d)_{J\to I}}{-\sum_{i_t} p(i_t)\log(p(i_t))} \quad (3)$$

Computing (normalized) TE in this way between all pairs of binned neuronal time series results in a time-scale dependent, weighted, directed network. Networks are weighted because some pairs of neurons fire more frequently and reliably at certain delays than others, and they are directed because a predictive, statistical relationship that exists from neuron J to neuron I, may not exist from neuron I to neuron J. Each element $a_{ij}$ in the TE matrix is the TE value from the $i^{th}$ to the $j^{th}$ neuron. TE values of zero denote the absence of an effective connection between the two neurons, while TE values greater than zero represent the weighted strength of the effective connection between the two neurons.

To determine the significance of network connections (TE values), TE values were computed for 5000 pairs of jittered spike trains. TE values which were larger than 99.9% of jittered values were considered significant, corresponding to a p-value of less than 0.001. Computing significant, normalized TE values for 25 recordings at two timescales, resulted in 50 full networks.

#### Quantification of neural computation

Computation by neurons receiving inputs from two other neurons in these networks was quantified following the Partial Information Decomposition (PID) from Williams and Beer (2010). The PID allows multivariate TE to be separated into distinct information components, one of which is a measure of neural computation termed synergy. The general form of the decomposition of multivariate TE between three neuronal time series, with two transmitter neurons, J and K, each sending a single input to one receiver neuron, I, can be expressed as (Figure 12):

$$\begin{gathered} TE(\{J,K\}\to I) = \text{Synergy}(\{J,K\}\to I) + \text{Unique}(K;J\to I) + \\ \text{Unique}(J;K\to I) + \text{Redundancy}(\{J,K\}\to I) \end{gathered} \quad (4)$$

where $\{J,K\}$ is a vector of the combined J and K time series. Similarly, we can express the decomposition of bivariate TE from neuron J to I and neuron K to I as (Figure 12):

$$TE(J\to I) = \text{Unique}(K;J\to I) + \text{Redundancy}(\{J,K\}\to I) \quad (5)$$

and

$$TE(K\to I) = \text{Unique}(J;K\to I) + \text{Redundancy}(\{J,K\}\to I) \quad (6)$$

In Equations 4-6, all terms are quantified in units of bits (see Williams and Beer 2010, 2011 for a full description of these terms). The unique terms correspond to the information provided by that time series alone (either the J or the K time series) about the current state of I. The redundant term represents the overlapping information provided by time series J and K about the current state of I. Notice, in Equations 5 and 6, that although TE is only dependent on the two time series that are directly interacting (either J and I, or K and I), because J and K are both interacting with the same time series, their unique interactions are influenced by each other. Thus, the unique information provided by one of these time series is dependent on the other. In other words, because J and K provide some redundant (overlapping) information about I, J influences how much information K provides uniquely versus redundantly about I. Likewise, K influences how much information J provides uniquely versus redundantly about I.

The synergistic term in Equation 4 is the additional information (beyond the unique and redundant information) that is processed by the receiver (I) based on the non-overlapping information from both inputs (J and K) occurring simultaneously. Thus, synergy is a proxy for the non-linear computation which takes information from two sources and combines them in some way to generate a unique output.

To calculate synergy, note that Equation 4 can be rewritten as:

$$\mathrm{Synergy}(\{J,K\} \to I) = TE(\{J,K\} \to I) - TE(J \to I) - TE(K \to I) + \mathrm{Redundancy}(\{J,K\} \to I) \tag{7}$$

by substituting Equations 5 and 6 and solving for synergy. Notice that we can compute all TE terms in Equation 7 via Equation 1. This leaves only the Redundancy term to be computed. Fortunately, a method for measuring this term has been provided by Williams and Beer (2010, 2011), who define redundancy in terms of a quantity titled the minimum information $I_{\min}$:

$$\begin{aligned}\mathrm{Redundancy}(\{J,K\} \to I) &\overset{\mathrm{def}}{=} I_{\min}(I_t; J_{t-1}K_{t-1} | I_{t-1}) = \\ &\sum_{i_t} p(i_t) \min_{R \in \{J_{t-1}, K_{t-1}\}} I_{spec}(I_t = i_t; R | I_{t-1}) = \\ &\sum_{i_t} p(i_t) \min_{R \in \{J_{t-1}, K_{t-1}\}} [I_{spec}(I_t = i_t; R, I_{t-1}) - I_{spec}(I_t = i_t; I_{t-1})]\end{aligned} \tag{8}$$

where the specific information $I_{spec}$ is defined as:

$$I_{spec}(I_t = i_t; R, I_{t-1}) = \sum_{r,\, i_{t-1}} p(r, i_{t-1} | i_t) \log\left(\frac{p(r, i_{t-1}, i_t)}{p(r, i_{t-1}) p(i_t)}\right) \tag{9}$$

and

$$I_{spec}(I_t = i_t; I_{t-1}) = \sum_{i_{t-1}} p(i_{t-1}|i_t) \log\left(\frac{p(i_{t-1}, i_t)}{p(i_{t-1})p(i_t)}\right) \tag{10}$$

Thus, redundancy is the minimum information provided by J or K about each state of I, averaged over all possible states. In other words, redundancy is the minimum overlapping information (the shared information) that the past states of J and K provide about the current states of I. Redundancy was calculated via Equations 9 and 10. Finally, synergy was calculated via Equation 7. Computing synergy for all possible triads (for each neuron that received at least two inputs, all possible groupings of two input neurons and the receiver were considered) in all networks yields a single synergy value per triad. We then normalized synergy values by dividing by the entropy of the future state of I, as done in Equation 3.

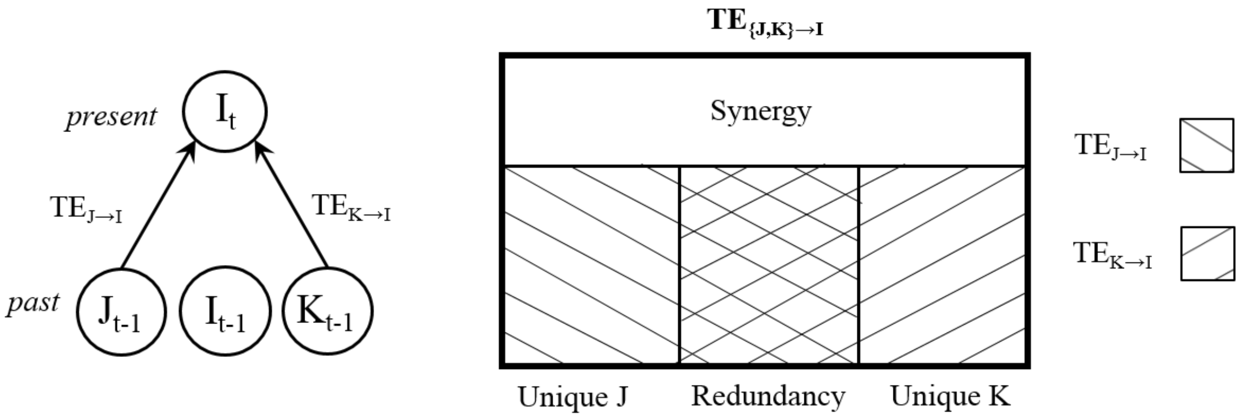


**Figure 12.** *The Partial Information Decomposition.* In this study, we analyzed two-input computations which were determined using the Partial Information Decomposition to dissect multivariate transfer entropy (occurring among three neurons, with two transmitter neurons each sending significant information to a receiver neuron) into synergistic, redundant, and unique information terms. The synergistic information component was used to represent the amount of computation carried out by the receiver.

Although there are other methods for calculating synergy (Bertschinger et al., 2014; Pica et al., 2017), we chose this measure because it is capable of detecting linear and nonlinear interactions and it is currently the only measure which can quantify how much synergy occurs in an interaction in which three variables (here, receiver past and pasts of the two transmitters) predict a fourth. Note, we chose not to consider higher order synergy terms, for systems with more than two transmitting neurons, due to the increased computational burden it presented (the number of PID terms increases rapidly as the number of variables increases). However, based on bounds calculated for the highest order synergy term by Timme et al. (2016), it was determined that the information gained by including an additional input beyond two either remained constant or decreased. Thus, it was inferred that lower order (two-input) computations dominated.

It is important to re-emphasize two things here. First, transmitter neurons and receiver neurons differ in terms of how they are defined. Transmitter neurons are required to have at least one outgoing connection (but not necessarily any incoming connections) and receiver neurons are required to have at least two incoming connections (but not necessarily any outgoing connections). Second, synergy occurs at the receiver neuron (where the two input signals are integrated). Thus, transmitters can be thought of as contributing to computation which occurs at the receiver neuron.

**Rich club detection**

To examine the relationship between rich clubs and computation in our networks, we identified the weighted rich clubs in each recording. Weighted rich clubs were identified using a modified version of the rich_club_wd.m function from the Matlab Brain Connectivity toolbox (Rubinov and Sporns, 2010; van den Heuvel and Sporns, 2011), adapted according to Opsahl et al. (2008) to compute weighted rich clubs. Briefly, this algorithm computes weighted rich-club coefficients as follows. For a given recording, a richness parameter ($r$), defined as the sum of the weights (incoming and outgoing), was computed for all neurons. Then, for every value of $r$ observed in a recording, a weighted rich club coefficient is computed. A rich club coefficient is computed for the $k^{\text{th}}$ value of $r$ as follows:

$$\Phi^{w}(r_k) = \frac{\sum_{i_{r \geq r_k}} \sum_{j_{r \geq r_k}} TE_{\{i,j\}}}{\sum^{n} TE^{rank}} \quad (6)$$

Here, the numerator is the amount of information transfer between neurons with $r$ greater than or equal to $r_k$, computed as the sum of the TE value between these neurons. The denominator is the maximum amount of information transfer that could have been observed among the neurons with $r$ greater than or equal to $r_k$. This is computed as the sum of the largest $n$ weights in the network where $n$ is the number of edges found between neurons with $r$ greater than or equal to $r_k$. The resulting ratio approaches one when the strongest connections connect the neurons that transfer the most information (i.e., richest neurons).

To establish the existence of a significant rich club at a given threshold $r_k$, we computed the ratio between the observed $\Phi^{w}(r_k)$ and the distribution of those observed when the edges of the network were shuffled. Shuffles were performed according to the methods of Maslov and Sneppen (2002). Briefly, this method randomly selects two edges (e.g., A→B and C→D) and randomly swaps either the sender or the receivers of the edges (A→D and C→B). Such rewiring only takes place if the newly created edges did not already exist in the network. To shuffle a network, the swapping process is repeated four times the number of edges in the network.

For each network, the rich club coefficients from shuffled versions of the network ($\Phi^{w}_{shuffled}$) were computed from 500 shuffled variants of the network. The mean coefficient for each threshold $r_k$ across shuffles was then used to standardize the observed coefficients, as follows:

$$\Phi^{w}_{std}(r_k) = \frac{\Phi^{w}_{observed}(r_k)}{|\Phi^{w}_{shuffled}(r_k)|} \quad (7)$$

The resulting standardized coefficient, $\Phi^{w}_{std}$, reflects how many times greater the observed coefficients are than the expected values given the distribution of edge weights observed in a given network. The significance of a given standardized coefficient was established by computing the associated p-value as the number of coefficients from shuffled networks that exceeded the observed coefficient and dividing by the number of shuffles (500). We defined p-value $\leq 0.01$ as significant.

#### Quantification of the synergy-rich club relationship

The relationship between synergy and rich clubs was performed using two approaches. In the first approach, we compared the amount of computation inside versus outside of the rich club by randomly selecting a single, significant representative rich club for each network and asking what the expected synergy-per-triad was for triads with receivers inside versus outside of the rich club. Given the risk for sampling bias introduced by the selection of representative rich clubs in the first approach, we also pursued a second approach that quantified the relationship between synergy and the rich club, at all possible thresholds. That is, for each network, we sorted neurons from strongest to weakest, and then cumulatively recruited neurons into the "rich club" one at a time. For this second approach, we computed the amount of computation inside and outside of the rich clubs defined at all possible thresholds for each network. The results were then aggregated across networks by aligning coefficients based on the percentage of the network included in the rich club. The above approaches were also used to compare the computation ratio, or the ratio of computation to propagation (summed triad TE), inside versus outside the rich club.

In addition to the above analyses which considered the position of the receiver with respect to the rich club, we also calculated the expected synergy found in all possible interactions of triads with respect to the rich club for all networks (see Figure 13). This was done to achieve a more detailed understanding of the relationship between triads, synergy, and rich clubs.

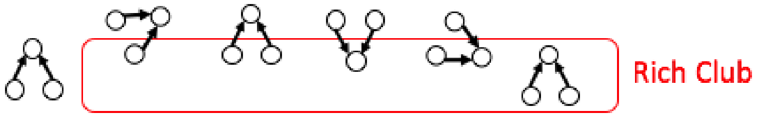


**Figure 13**. Schematic of possible configurations of synergistic triads interacting with the rich club. In order to quantify the amount of computation that takes place within the rich club and to determine how the amount of computation depends on the interaction between synergistic triads and the rich club, we considered all ways in which synergistic triads could interact with the rich club. From left to right these include: no triad nodes or edges participate in the rich club, a single transmitter (arrow pointing away) is in the rich club, both transmitters are in the rich club, only the receiver (both arrows pointing toward) is in the rich club, a transmitter-edge-receiver combination is in the rich club, and finally, the entire connected triad is in the rich club.

#### Alternative approach to PID

Because there are many methods for determining whether or not a neuron computes based on its inputs, we chose to consider an alternative approach to PID which calculates neuron transfer functions according to the method proposed by Chichilnisky (2001). This method begins with the calculation of each neuron's spike-triggered average (STA), which is the average pattern of input spikes preceding a spike in the cell (i.e. the sum of the inputs $i$ preceding each spike, divided by the total number of spikes $f$) for a specific delay $d$:

$$\mathrm{STA} = \frac{\sum_{t=1}^{d} i_t f_t}{\sum_{t=1}^{d} f_t} \quad (8)$$

Here, we considered a delay of 1-14 ms. In other words, we looked at input patterns occurring anywhere from 14 to 1 ms before the spike of each neuron. To determine each neuron's response based on its STA, we next calculated a *generator signal*: $g_t = a \cdot s_t$, which is a linear combination of the input spikes at a particular time, and then examined the average spike count in time bins with approximately equivalent generator signals (Chichilnisky, 2001). This gave us a neuron transfer function in the form of probability of spiking vs. generator signal (number of inputs). Because this relationship can be linear or nonlinear, we then fit each neuron's transfer function with both a linear and nonlinear (sigmoidal) fit, and calculated the sum of squared errors for each fit. We then computed the ratio of the sum of squared errors for the sigmoidal fit to sum of squared error for the linear fit and used a median split to classify neurons as having either linear or nonlinear transfer functions. Neurons whose sum of squared errors ratio was greater than the median were classified as linear, whereas those whose sum of squared errors ratio was less than the median were classified as nonlinear.

To examine how this approach relates to PID, we compared the expected synergy for neurons with nonlinear versus linear transfer functions across all networks. To parallel our synergy-rich club analysis, we also compared the concentration of neurons with nonlinear transfer functions inside and outside the rich club. That is, we computed the percentage of rich club and non-rich club neurons that had nonlinear transfer functions.
